# AlzGPS: A Genome-wide Positioning Systems Platform to Catalyze Multi-omics for Alzheimer’s Therapeutic Discovery

**DOI:** 10.1101/2020.09.17.302612

**Authors:** Yadi Zhou, Jiansong Fang, Lynn Bekris, Young Heon Kim, Andrew A. Pieper, James B. Leverenz, Jeffrey Cummings, Feixiong Cheng

**Affiliations:** Genomic Medicine Institute, Lerner Research Institute, Cleveland Clinic, Cleveland, OH 44195, USA; Department of Molecular Medicine, Cleveland Clinic Lerner College of Medicine, Case Western Reserve University, Cleveland, OH 44195, USA; Harrington Discovery Institute, University Hospitals Cleveland Medical Center, Cleveland, OH 44106, USA; Department of Psychiatry, Case Western Reserve University, Cleveland, OH 44106, USA; Geriatric Psychiatry, GRECC, Louis Stokes Cleveland VA Medical Center; Cleveland, OH 44106, USA; Institute for Transformative Molecular Medicine, School of Medicine, Case Western Reserve University, Cleveland 44106, OH, USA; Weill Cornell Autism Research Program, Weill Cornell Medicine of Cornell University, New York, NY 10065, USA; Department of Neuroscience, Case Western Reserve University, School of Medicine, Cleveland, OH 44106, USA; Lou Ruvo Center for Brain Health, Neurological Institute, Cleveland Clinic, Cleveland, Ohio 44195, USA; Cleveland Clinic Lou Ruvo Center for Brain Health, Las Vegas, NV 89106, USA; Chambers-Grundy Center for Transformative Neuroscience, Department of Brain Health, School of Integrated Health Sciences, UNLV, Las Vegas, Nevada 89154, USA; Case Comprehensive Cancer Center, Case Western Reserve University School of Medicine, Cleveland, OH 44106, USA

**Keywords:** Alzheimer’s disease, network medicine, database, drug repurposing, clinical trial, transcriptomics

## Abstract

**Background:** Over15 million family members and caregivers have expended $220 billion for care of patients with AD and other dementias, and the attrition rate for AD clinical trials (2002-2012) is estimated at 99.6%. While recent DNA/RNA sequencing and other multi-omics technologies have advanced the understanding of the biology and pathophysiology of AD, no effective disease-modifying or preventive therapies, for AD have emerged in the past two decades. A new approach to integration of the genome, transcriptome, proteome, and human interactome in the drug discovery and development process is essential for this endeavor.

**Methods:** In this study, we developed AlzGPS (<u>G</u>enome-wide <u>P</u>ositioning <u>S</u>ystems platform for <u>Alz</u>heimer’s Therapeutic Discovery, https://alzgps.lerner.ccf.org), a comprehensive systems biology tool to enable searching, visualizing, and analyzing multi-omics, various types of heterogeneous biological networks, and clinical databases for target identification and effective prevention and treatment of AD.

**Results:** Via AlzGPS: (1) we curated more than 100 AD multi-omics data sets capturing DNA, RNA, protein, and small molecules’ profiles underlying AD pathogenesis (e.g., early vs. late stage and tau vs. amyloid endophenotype); (2) we constructed endophenotype disease modules by incorporating multi-omics findings and human protein-protein interactome networks; (3) we identified repurposable drugs from ∼3,000 FDA approved/investigational drugs for AD using state-of-the-art network proximity analyses; (4) we curated 300 literature references for highly repurposable drugs; (5) we included information from over 200 ongoing AD clinicals noting drug mechanisms and primary drug targets, and linking them to our integrated multi-omics view for targets and network analyses results for the drugs; (6) we implemented a highly interactive web-interface for database browsing and network visualization.

**Conclusions:** Network visualization enabled by the AlzGPS includes brain-specific neighborhood networks for genes-of-interest, endophenotype disease module networks for data sets-of-interest, and mechanism-of-action networks for drugs targeting disease modules. By virtue of combining systems pharmacology and network-based integrative analysis of multi-omics data, the AlzGPS offers actionable systems biology tools for accelerating therapeutic development in AD.

## Background

Alzheimer’s disease (AD) is a progressive neurodegenerative disorder accounting for 60-80% of dementia cases (1). In addition to cognitive decline, AD patients have extensive neuropathological changes including deposition of extracellular amyloid plaques, intracellular neurofibrillary tangles, and neuronal death (2, 3). It is estimated that the number of AD patients will reach 16 million by 2050 in the United States alone (4, 5). Effective treatments are needed, as there are no disease-modifying treatments for AD and no new drugs have been approved since 2003 by the US Food and Drug Administration (FDA). There are several possible explanations for the high failure rate in AD drug discovery. For example, transgenic rodent models used to test drugs may not fully represent human AD pathobiology (6). Also, there is a lack of sensitive measures for outcomes in clinical trials. Other potential immediate causes for clinical trial failures include targeting the wrong pathobiological or pathophysiological mechanisms, attempted intervention at the wrong stage (too early or too late), unfavorable pharmacodynamic and pharmacokinetic characteristics of the drug (e.g., poor brain penetration), lack of target engagement by drug candidates, and hypothesis that fail to incorporate the great complexity of AD (6, 7).

Multiple types of omics data have greatly facilitated our understanding of the pathobiology of AD. For example, using single-cell RNA-seq, a novel microglia type (termed disease-associated microglia, DAM) was discovered to be associated with AD, understanding of whose molecular mechanism could offer new therapeutic targets (8). Using large-scale genome-wide association studies (GWAS), twenty loci showed genome-wide significant association with Alzheimer’s disease, among which 11 were newly discovered (9). A recent study using deep profiling of proteome and phosphoproteome prioritized proteins and pathways associated with AD, and it was shown that protein changes and their corresponding RNA levels only partially coincide (10). The large amount of multi-omics data and recent advances in network-based methodologies for drug repurposing today present unprecedented opportunities for accelerating target identification for drug discovery for AD, and this potential has also been demonstrated in other complex diseases as well, such as cancer (11), cardiovascular disease (12), and schizophrenia (13), and are beginning to be exploited in AD (6, 14). Drug repurposing offers a rapid and cost-effective solution for drug discovery for complex disease, such as the current global pandemic of coronavirus disease 2019 (COVID-19) (15, 16) and AD (6). The central idea of network-based drug repurposing is that for a drug to be able to affect a disease, the drug targets must directly overlap with or be in the immediate vicinity of the disease modules, which can be identified using the vast amount of high-throughput sequencing data (**Figure 1A**). Our recent efforts using network-based methodologies and AD omics data have led to the discovery of two drugs that show efficacy in network models in AD: sildenafil and pioglitazone (14). Network analysis provides potential mechanisms for these drugs and facilitates experimental validation. Therefore, we believe posit that a comprehensive systems biology tool in the framework of network-based multi-omics analysis could inform Alzheimer’s patient care and therapeutic development.

**Figure 1.**
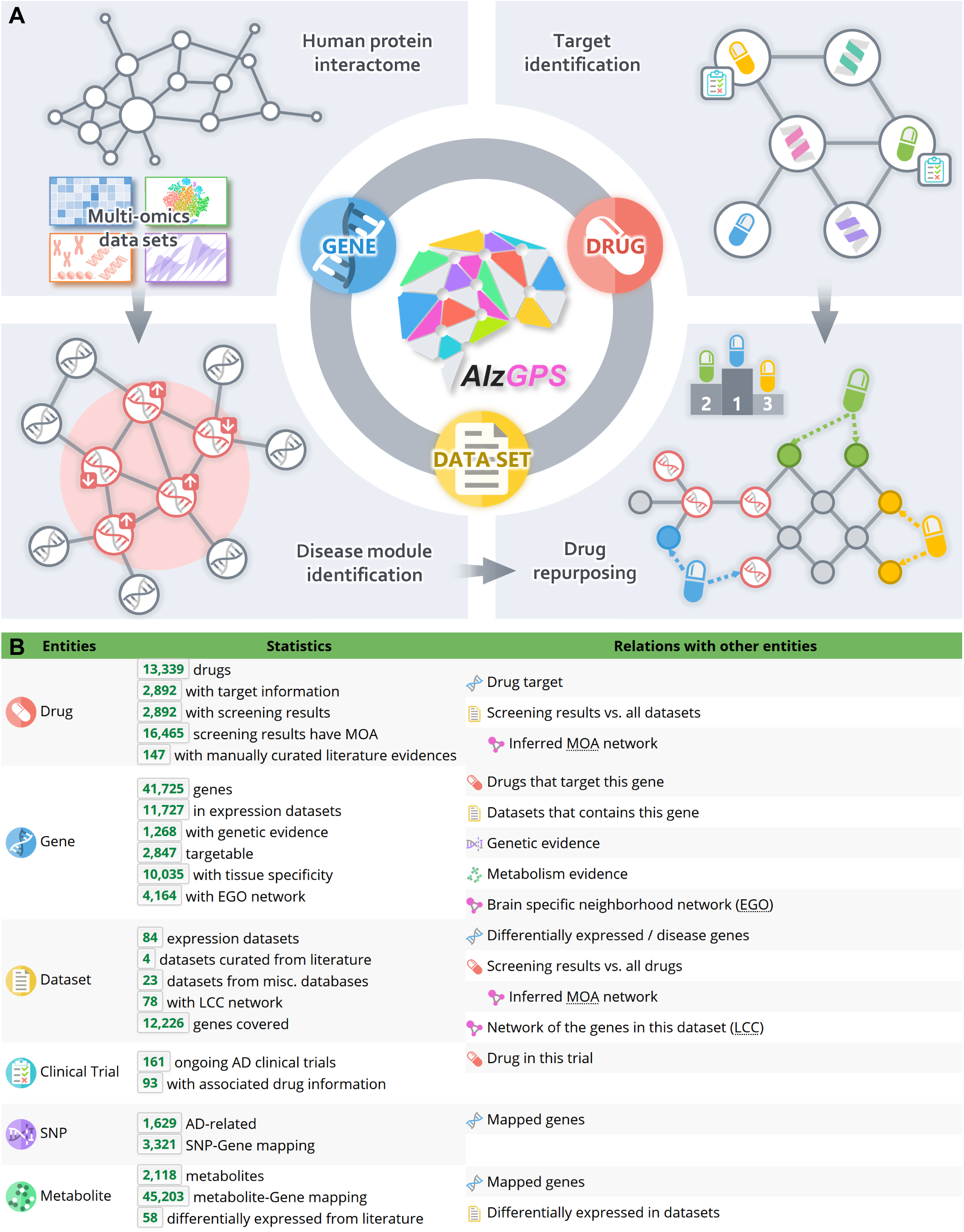
The architecture of AlzGPS. (**A**) AlzGPS is built on three main data entities (genes, drugs, and omics layers) and their relationships. The multi-omics data (genomics, transcriptomics (bulk and single-cell/single-nucleus, and proteomics) in AlzGPS help identify likely causal genes/targets that are associated with Alzheimer’s disease (AD) and disease modules in the context of human protein-protein interactome. Then, via network proximity measure between drug-target networks and disease modules in the human interactome, drugs can be prioritized for the potential to alter the genes in the module, for potential treatment of AD. (**B**) Detailed statistics of the entities and relations in AlzGPS. EGO – brain-specific neighborhood network (ego network); LCC – largest connected component network; MOA – mechanism-of-action network.

To this end, we present a new freely-available database and tool, named AlzGPS (A <u>G</u>enome-wide <u>P</u>ositioning <u>S</u>ystems platform for <u>Alz</u>heimer’s Therapeutic Discovery), for target identification and drug repurposing for AD. AlzGPS was built with large scale diverse information, including multi-omics (genomics, (bulk and single cell) transcriptomics, proteomics, and interactomics) of human and other species, drug-target network, literature-derived evidence, AD clinical trials information, and network proximity analysis (**Figure 1B**). Our hope is that AlzGPS will be a valuable resource for the AD research community for several reasons. First, AlzGPS contains abundant multi-domain information types all coalesced in one location. The manually curated data, such as the literaturederived information for the most promising repurposable drugs and more than 100 multi-omics AD data sets, are of high quality and relevance. Second, using state-of-the-art network proximity approaches, AlzGPS provides a systemic evaluation of 3000 FDA approved or investigational drugs against the AD data sets. These results (along with various network visualizations) will provide insights for potential repurposable drugs with clear network-based footprints in the context of the human protein interactome. The drug-data set associations can be further explored in AlzGPS for individual drug targets or genes associated with AD. Lastly, AlzGPS offers a highly interactive and intuitive modern web interface. The relational nature of these data was embedded in the design to help the user easily navigate through different types of information. In addition, AlzGPS provides three types of network visualizations for the tens of thousands of networks in the database, including brain-specific neighbor networks for genes, disease modules for data sets, and inferred mechanism-of-action (MOA) networks for drugs and data set pairs with significant proximity. AlzGPS is freely available to the public without registration requirement at https://alzgps.lerner.ccf.org.

## Methods

### Data collection and preprocessing

#### AD data sets

A data set is defined as either (1) genes/proteins/metabolites that are differentially expressed in AD patients/mice versus controls; or (2) genes that have known associations with risks of AD from literature or other databases. We retrieved expression data sets underlying AD pathogenesis capturing transcriptomics (microarray, bulk or single-cell RNA-Seq) and proteomics across human, mouse, and model organisms (e.g. fruit fly and *C. elegans*). All the samples of the data sets were derived from total brain, specific brain regions (including hippocampus, cortex, and cerebellum), and brain-derived single cells, such as microglial cells. For some of the expression data sets, the differentially expressed genes/proteins were obtained from the original publications (from main tables or supplemental tables). For other data sets that did not have such differential expression results available, the original brain microarray/RNA-Seq data were obtained from Gene Expression Omnibus (GEO) (17) and differential expression analysis was performed using the tool GEO2R (18). GEO2R performs the differential expression analysis for the sample groups defined by the user using the limma R package (19). All differentially expressed genes identified in mouse were further mapped to unique human-orthologous genes using the NCBI HomoloGene database (https://www.ncbi.nlm.nih.gov/homologene). The details for all the data sets, including organism, genetic model (for mouse), brain region, cell type (for single-cell RNA-Seq), PubMed ID, GEO ID, and the sources (e.g., supplemental table or GEO2R), etc., can be found in Table S1.

#### Genes and Proteins

We retrieved the gene information from the HUGO Gene Nomenclature Committee (HGNC, https://www.genenames.org/) (20), including gene symbol, name, type (e.g., coding and non-coding), chromosome, synonyms, and identification (ID) mapping in various other databases such as National Center for Biotechnology Information (NCBI) Gene, ENSEMBL, and UniProt. All proteins from the AD proteomics data sets were mapped to genes using the mapping information from HGNC.

#### Single-nucleotide polymorphisms (SNPs)

We found 3,321 AD-associated genetic records for 1,268 genes mapped to 1,629 SNPs, by combining results from GWAS Catalog (https://www.ebi.ac.uk/gwas/) (21) using the trait “Alzheimer’s disease” and published studies. The PubMed IDs for the genetic evidence are provided on AlzGPS.

#### Tissue expression specificity

We downloaded RNA-Seq data (Reads Per Kilobase Million [RPKM] value) of 32 tissues from GTEx V6 release (accessed on April 01, 2016, https://gtexportal.org/home/). We defined the genes with RPKM≥1 in over 80% of samples as tissue-expressed genes and the other genes as tissue-unexpressed. To quantify the expression significance of tissue-expressed gene *i* in tissue *t*, we calculated the average expression <*E*(*i*)> and the standard deviation *δ_E_*(*i*) of a gene’s expression across all included tissues. The significance of gene expression in tissue *t* is defined as

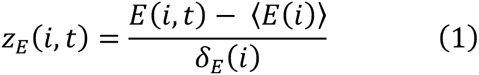

#### Drugs

We retrieved drug information from the DrugBank database (v4.3) (22), including name, type, group (approved, investigational, etc.), Simplified Molecular-Input Line Entry System (SMILES) and Anatomical Therapeutic Chemical (ATC) code(s). We also evaluated the pharmacokinetic properties (such as blood–brain barrier [BBB] penetration) of the drugs using admetSAR (23, 24).

#### Drug literature information for AD treatment

For the top 300 repurposable drugs (i.e., drugs with the highest number of significant proximities to the AD data sets), we manually searched and curated the literature for their therapeutic efficacy against AD using PubMed. In addition to the title, journal, and PubMed ID, we summarized the types (clinical and non-clinical), experimental settings (e.g., mouse/human and transgenic line for non-clinical studies; patient groups, randomization type, length, and control type of clinical studies), and results of these studies. In total, we found 292 studies for 147 drugs.

#### Drug-target network

To build a high-quality drug-target network, several databases were accessed, including the DrugBank database (v4.3) (22), Therapeutic Target Database (TTD) (25), PharmGKB database, ChEMBL (v20) (26), BindingDB (27), and IUPHAR/BPS Guide to PHARMACOLOGY (28). Only biophysical drug-target interactions involving human proteins were included. To ensure data quality, we kept only interactions that have inhibition constant/potency (K_i_), dissociation constant (K_d_), median effective concentration (EC_50_), or median inhibitory concentration (IC_50_) ≤ 10 µM. The final drug-target network contains 21,965 interactions among 2,892 drugs and 2,847 human genes.

#### Clinical trials

The AD intervention clinical trials were retrieved from Cummings *et al.* 2018 (29) & 2019 (30). Information including phase, posted date, status, and agent(s) was obtained from https://clinicaltrials.gov. Drugs were mapped to the DrugBank IDs. Proposed mechanism and therapeutic purpose were from Cummings *et al.* 2018 (29) & 2019 (30).

#### Human protein interactome

We used our previously built high-quality comprehensive human protein interactome which contains 351,444 unique protein-protein interactions (PPIs, edges) among 17,706 proteins (nodes) (11, 12, 31, 32). Briefly, five types of evidence were considered for building the interactome: physical PPIs from protein three-dimensional (3D) structures, binary PPIs revealed by high-throughput yeast-two-hybrid (Y2H) systems, kinase-substrate interactions by literature-derived low-throughput or high-throughput experiments, signaling networks by literature-derived low-throughput experiments, and literature-curated PPIs identified by affinity purification followed by mass spectrometry (AP-MS), Y2H, or by literature-derived low-throughput experiments. No inferred PPIs were included.

### Network proximity quantification of drugs and AD data sets

To quantify the associations between drugs and AD-related gene sets from the data sets, we adopted the “closest” network proximity measure:

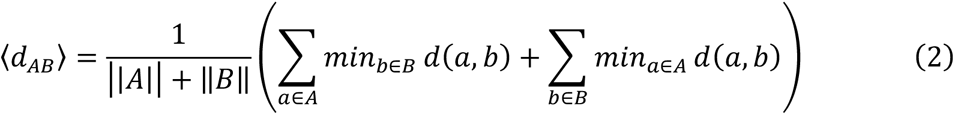

where *d*(*a*, *b*) is the shortest path length between gene *a* and *b* from gene list *A* (drug targets) and *B* (AD genes), respectively. To evaluate whether such proximity was significant, we performed z score normalization using a permutation test of 1,000 repeats. In each repeat, two randomly generated gene lists that have similar degree distributions to A and *B* were measure for the proximity. The z score was calculated as:

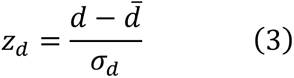

P value was calculated according the permutation test. Drug-data set pairs with Z < -1.5 and P < 0.05 were considered significantly proximal. In addition to network proximity, we calculated two additional metrics, overlap coefficient *C* and Jaccard index *J*, to quantify the overlap and similarity of *A* and *B*:

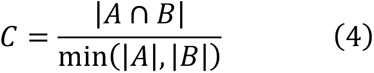

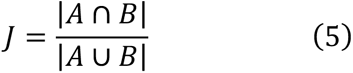

### Generation of networks

We offer three types of networks on AlzGPS: brain-specific neighborhood (EGO) network for the genes, largest connected component (LCC) network for the data sets, and inferred MOA network for significantly proximal drug-data set pairs. The three networks differ by inclusion criteria of the nodes (genes/proteins). The edges are PPIs colored by their types (e.g., 3D, Y2H, and literature). All networks are colored by whether they can be targeted by the drugs in our database.

For the EGO networks, we filtered genes by their brain expression specificity and generated only the network for those with positive brain specificity. We used the *ego_graph* function from NetworkX (33) to generate the EGO networks. The networks are centered around the genes-of-interest. An LCC network was generated for each AD data set using the *subgraph* function from networkx. For MOA, we examined the connections (PPIs) among the drug targets and the data sets.

### Website implementation

AlzGPS was implemented with the Django v2.2.2 framework (www.djangoproject.com). The website frontend was implemented with HTML, CSS, and JavaScript. The frontend was designed to be highly interactive and integrative. It uses AJAX to asynchronously acquire data in JSON format based on user requests to dynamically update the frontend interface. This architecture can therefore be integrated into end users’ own pipelines. Network visualizations were implemented using Cytoscape.js (34).

## Results and Discussion

### Information architecture and statistics

One key feature of AlzGPS is the highly diverse yet interconnected data types (**Figure 1**). The three main data types are genes, drugs, and AD-relevant omics data sets. More than 100 omics data sets were processed, including 84 expression data sets (**Table S1**) from AD transgenic animal models or patient-derived samples and 27 data sets from the literature or acquired from other databases. The expression data sets contain transcriptomic and proteomic data of human and rodent samples. Comparative sample groups were available in these data sets, such as early stage vs. late stage, healthy vs. AD. The differentially expressed genes/proteins were calculated for each data set.

The statistics and relations of the database are shown in **Figure 1B**. We collected and processed all the basic information (see **Methods**) and then constructed the relationships among the data types. For example, for genes and drugs, the relationship is drugs targeting proteins (genes); for gene and data set, the relationship is genes being differentially expressed in the expression data sets or included in other types of data sets, such as literature-based; for drug and data set, the proximity between each pair was calculated (see **Methods**) to identify the drugs that are significantly proximal to a data set, and vice versa.

Additional data types were collected or generated. For genes, these included genetic evidence (variants associated with AD) and tissue expression specificity to provide additional information for target gene identification. For drugs, we collected the data from ongoing clinical trials, including the proposed mechanism and therapeutic purpose (29) & (30). The trials were mapped to drugs. The BBB probability was computed (23, 24). For the top 300 drugs with the highest number of significant proximities to all the data sets, we manually curated the available literature. A total of 292 studies were found for 147 drugs (49%) that reported the associations of the drugs and AD. We grouped these studies into clinical and non-clinical, and extracted trial information for clinical type and experimental setting (number and type of patients) for both types. We also summarized and provide the study results.

### Web interface and network visualizations

A highly interactive web interface was implemented (**Figure 2**). On the home page (**Figure 2A**), the user can search for drugs, genes, metabolites, and gene variants. The user can directly list all drugs by their first-level ATC code, all AD data sets available, and all the ongoing clinical trials (**Figure 2B**). The search results are displayed in the “DATA TABLE” tab and switched with their associated buttons in the “RESULT” section on the left. Each data entity has its own data table for the associated information in the “DATA TABLE” tab. For example, on the gene page of APP (**Figure 2B**) is the basic information (green rows), such as name, type, chromosome, and synonym; descriptions for the derived data (purple rows), such as tissue specificity and number of genetic records; and external links (red row). Data for the relations of APP and other entities can be loaded by clicking the button in “DETAIL” (blue row). For example, the expression data sets in which APP is differentially expressed can be found by clicking the “Dataset” button (**Figure 2B**). Any data loaded will be added to the same explorer. The buttons in the “RESULT” are organized in trees. For example, APP is included in the “V1 AD-seed” data set, which contains 144 AD-associated genes with strong literature evidence. When the user clicks this data set in the APP gene table, a new data table for the “V1 AD-seed” data set will replace the the APP gene page, and a new button with indentation will appear below the APP button in “RESULT” (**Figure 2B**).

**Figure 2.**
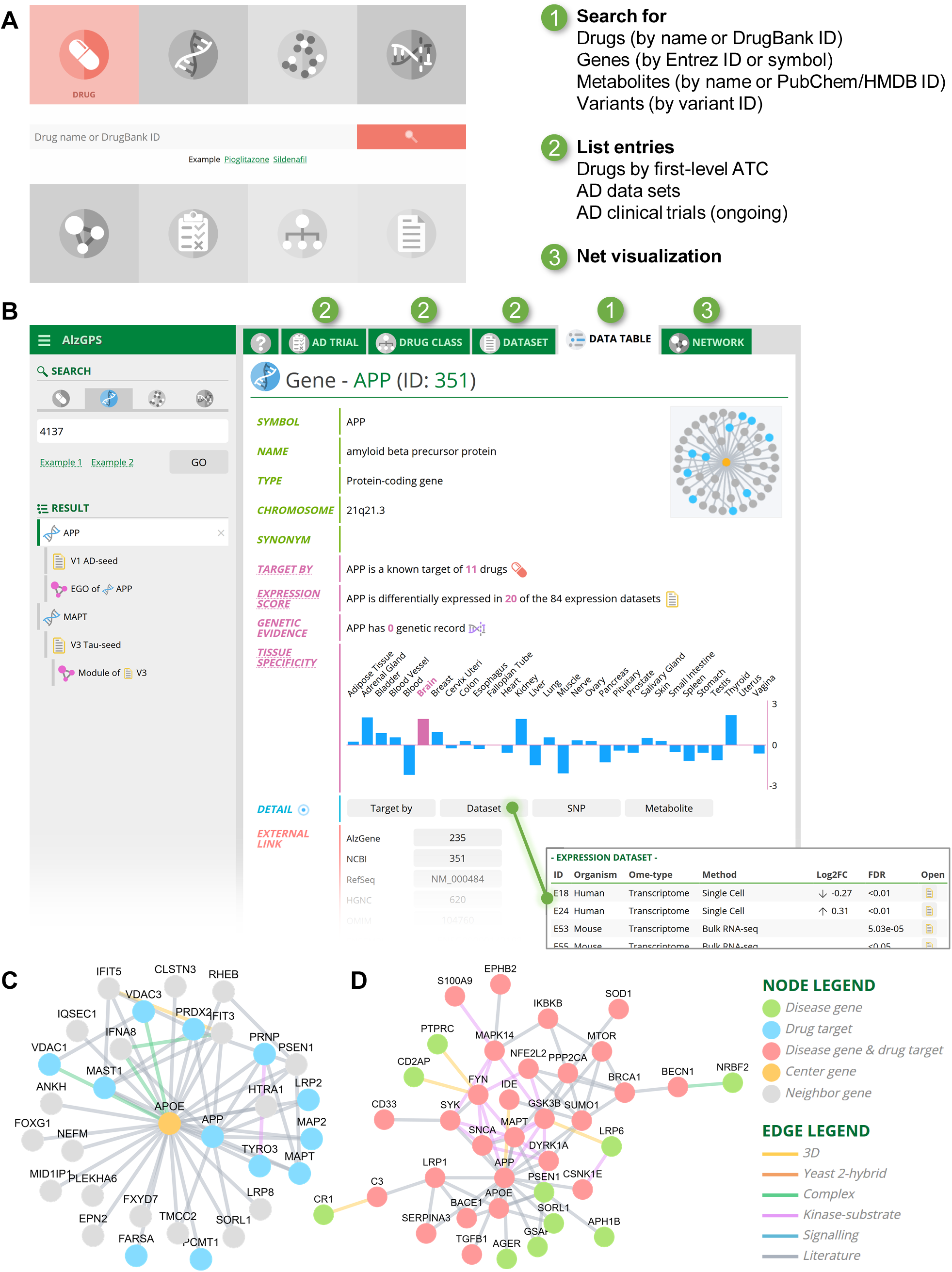
Web interface overview. (**A**) The home page provides access to searching, listing entries, and viewing brain-specific gene/target networks. User will be redirected to the interactive explorer (**B**), in which all information are then dynamically loaded and added to the same web page. Each data entity has its own basic information page under the “DATA TABLE” tab. Additional information regarding the relations (e.g., proximity results) can be loaded by clicking the corresponding button in the “DETAIL” section. (**C**) An example brain-specific neighborhood network using APOE. (**D**) An example largest connected component network using data set “V2”.

An all-in-one interactive explorer that minimizes the need for navigation of information using the relational nature of these data is a major feature of the web interface. Another major feature is the network visualizations. We offer three types of networks, (1) the brain-specific neighborhood network (EGO) for a gene-of-interest that shows the PPIs with its neighbors (**Figure 2C**); (2) the largest connected component (LCC) network for a data set that shows the largest module formed by the genes in this data set (**Figure 2D**); and (3) inferred MOA network for a significantly proximal drug-data set pair, which is illustrated in the case studies below.

### Case study – target identification

Generally, using AlzGPS for AD target identification starts with selecting one or a set of data sets (**Figure 2B**, “DATASET” tab). Users can select a data set based on organisms, methods (e.g., single-cell/nuclei RNA-Seq), brain regions, and comparisons (e.g., early-onset AD vs healthy control) for the expression data sets. Additionally, we have collected data sets from the literature, other databases, or computationally predicted results. Here, we use the “V1 AD-seed” data set as a starting point. This data set was from our recent study which contains 144 AD-associated genes based on literature-derived evidence. We found that 118 genes were differentially expressed as shown in at least one data set. By browsing these genes, we selected four examples, microtubule associated protein tau (MAPT), bridging integrator 1 (BIN1), apolipoprotein E (APOE), and β-secretase 1 (BACE1) based on positive brain expression specificity and number of data sets that include them.

#### MAPT

MAPT encodes the tau protein, modification of which is one of the main neuropathological hallmarks of AD (35, 36). Mutations and alternative splicing of MAPT are associated with risk of AD (37). MAPT is differentially expressed in five expression data sets (**Figure 3A**) and has high brain specificity. Five pieces of genetic evidence were found for MAPT. MAPT can be targeted by 27 drugs. In addition, many of its direct PPI neighbors are targetable, suggesting a potential treatment strategy by targeting MAPT and its neighbors.

**Figure 3.**
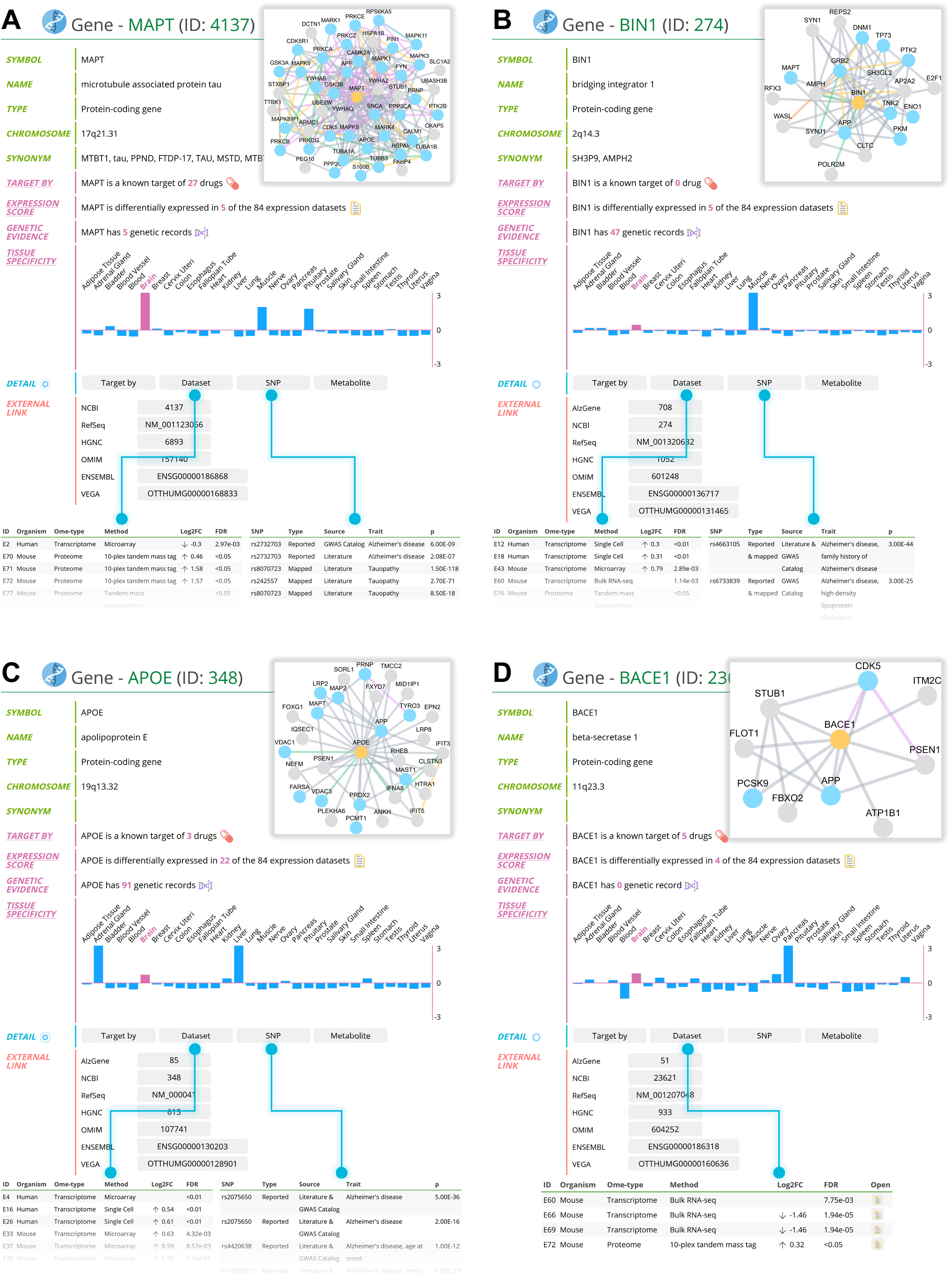
Case study – target identification. Four genes, MAPT (**A**), BIN1 (**B**), APOE (**C**), and BACE1 (**D**) are used as examples to show the gene page. On the gene page, we show a summary of several statistics of the gene in AlzGPS, including the number of drugs that can target it, number of data sets of omics in which the target/protein coding gene is differentially expressed, number of genetic records, and the brain-expression specificity. Detailed information can be loaded by clicking corresponding buttons. Examples of detailed differential expression results and genetic records are shown for these four genes. In addition, a brain-specific neighborhood network is available that centers around the gene-of-interest and show the targetability of its neighborhood.

#### BIN1

BIN1 is one of the most important susceptibility genes for late-onset AD (38), and can modulate tau pathology (39). Higher levels of BIN1 expression are associated with a delayed age of AD onset (40). Differentially manifested in five data sets, BIN1 has 47 genetic record associations (**Figure 3B**). Although no drugs are known to target BIN1, many of the BIN1’s PPI neighbors can be targeted.

#### APOE

The ε4 allele of APOE is the main genetic risk factor of AD (41). Apolipoprotein E ε4 plays an important role in Aβ deposition (41), a major pathological hallmark of AD. APOE is differentially expressed in 22 data sets (**Figure 3C**). It has a high number of associated genetic records – 91. Both APOE and its PPI partners can be targeted.

#### BACE1

β-secretase 1 (BACE1) cleaves APP and generates amyloid-β peptides (42), whose aggregation is another pathological hallmark of AD. The inhibition of BACE1 has been a popular target for AD drug development. Shown in **Figure 3D**, BACE1 is differentially expressed in 4 data sets.

### Case study – drug repurposing

In this section, we use sildenafil and pioglitazone as two examples. In our recent studies, we found that both sildenafil and pioglitazone were associated with a reduced risk of AD using network proximity analysis and retrospective case-control validation (14). Mechanistically, *in vitro* assays showed that both drugs were able to downregulate cyclin-dependent kinase 5 (CDK5) and glycogen synthase kinase 3 beta (GSK3B) in human microglia cells. These drugs were discovered using different data sets. Sildenafil was found using a high-quality literature-based AD endophenotype module (available as AlzGPS data set “V1 AD-seed”) containing 144 genes. Pioglitazone was found using 103 high-confidence AD risk genes (available as AlzGPS data set “V4 AD-inferred-GWAS-risk-genes”) identified by GWAS (13).

AlzGPS provides a list-view of the network proximity results of all the drugs organized by their first-level ATC code, which can be found in the “DRUG CLASS” tab (**Figure 2B**). The drugs are ranked by the number of significant proximities to the data sets. Sildenafil is the top four of the 148 drugs under the ATC code G “Genito-urinary system and sex hormones” with network proximity results, the top three being vardenafil, ibuprofen, and gentian violet cation. Pioglitazone is the top sixth of the 226 drugs under the ATC code A “Alimentary tract and metabolism”, following tetracycline, human insulin, epinephrine, cholecalciferol, and teduglutide. Both drugs achieved high numbers of significant proximities to the expression data set. Next, we examined the basic information of these drugs (**Figure 4A** and **4E**). Both drugs are predicted to be BBB penetrable. Sildenafil has 20 known targets and is significantly proximal to 27 of the 111 data sets (**Figure 4A**). We found one non-clinical study that reported that sildenafil treatment improves cognition and memory of vascular dementia in aged rats (43) (**Figure 4C**). As noted, we identified the potential of sildenafil against AD using the AD endophenotype module (**Figure 4B**, Z = -2.44, P = 0.003). Then, clicking the corresponding “MOA (mechanism-of-action)” button opened the inferred MOA network for sildenafil and the data set (**Figure 4D**). Although sildenafil does not target the genes in the data set (green) directly, it can potentially alter them through PPIs with its targets (blue).

**Figure 4.**
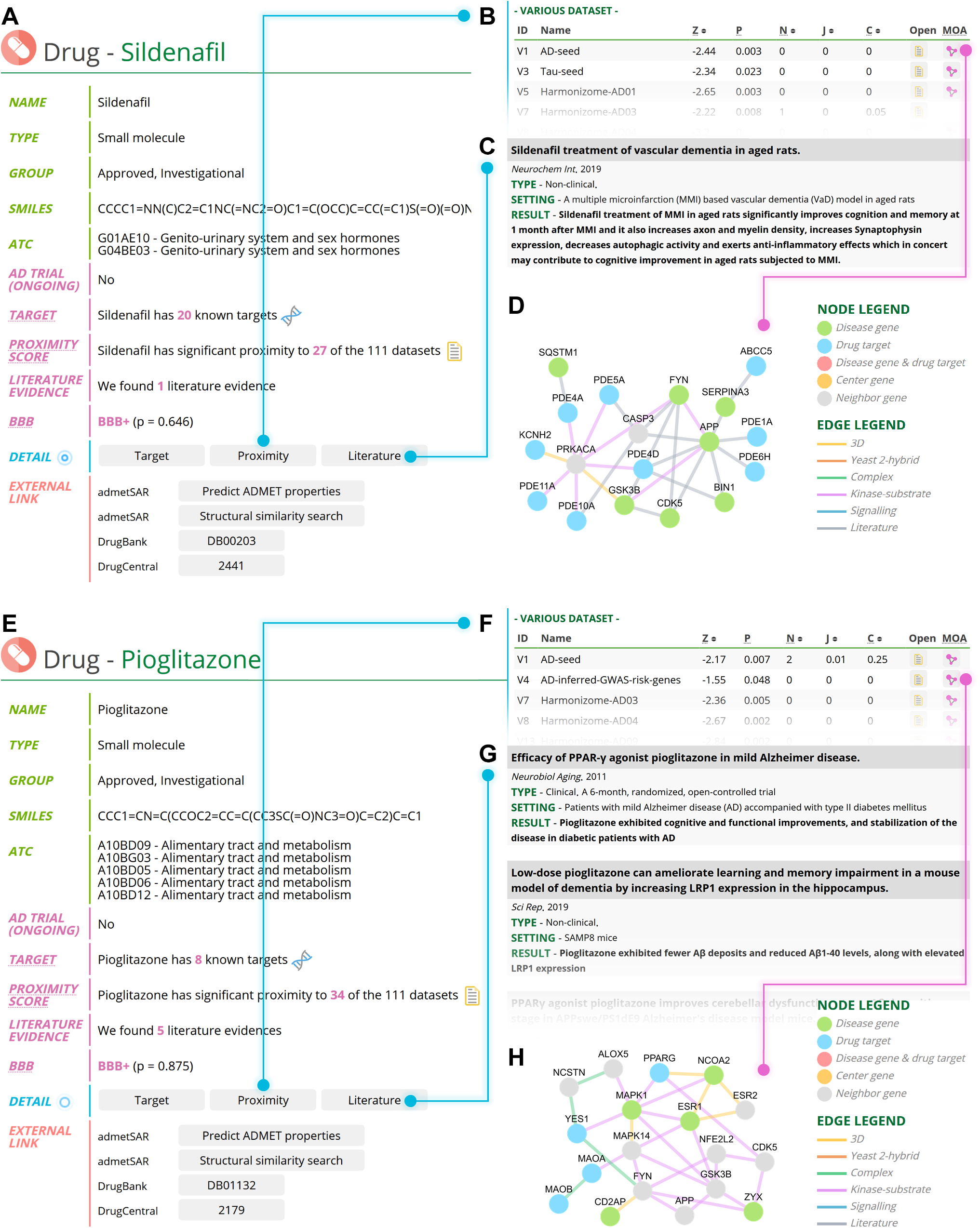
Case study – drug repurposing. Sildenafil and pioglitazone are used as examples to demonstrate how to use AlzGPS for drug repurposing. (**A**) Basic information for sildenafil. (**B**) Network proximity results for sildenafil. (**C**) Literature evidence for sildenafil. (**D**) Inferred mechanism-of-action for sildenafil targeting the “V1 AD-seed” data set, which contains 144 high-quality literature-based Alzheimer’s disease (AD) endophenotype genes. (**E**) Basic information for pioglitazone. (**F**) Network proximity results for pioglitazone. (**G**) Five studies were found that were related to treating AD with pioglitazone. (**H**) Inferred mechanism-of-action for pioglitazone targeting the “V4 AD-inferred-GWAS-risk-genes” data set which contains 103 high-confidence AD risk genes identified using genome-wide association studies.

Pioglitazone has 8 known targets and is significantly proximal to 34 data sets (**Figure 4E**). Five studies, containing both clinical and non-clinical data were found to be related to treating AD with pioglitazone. For example, a clinical study showed that pioglitazone can improve cognition in AD patients with type II diabetes (44) (**Figure 4G**). Similarly, network results and associated MOA networks suggested that pioglitazone can affect AD risk genes through PPIs (**Figure 4F** and **Figure 4H**).

### Validation studies

Once candidate agents are identified on AlzGPS, a variety of validation steps can be pursued (6). The agent can be tested in animal model systems of AD pathology to evaluate the predicted MOA of behavioral and biological effects. Since these are repurposed agents and have been used for other indications in human healthcare, electronic medical records can be interrogated to determine if there are notable effects on AD incidence, prevalence, or rate of progression. Both these methods are imperfect since animal models have rarely been predictive of human response, and doses and duration of exposures may be different for indications of other then AD in which the candidate agents are used. The ultimate assessment that could make an agent available for human care is success in a clinical trial and nominated agents must eventually be submitted to trials. If repurposed agents are not entered into trials because of intellectual property limitations or other challenges, the information from AlzGPS may be useful in identifying druggable disease pathways or providing seed structures that provide a basis for creation of related novel agents with similar MOAs.

## Conclusions

AlzGPS contains rich and diverse information connecting genes, AD data sets, and drugs for AD target identification and drug repurposing. It utilizes multiple biological networks and omics data such as genomics, transcriptomics, and proteomics, and provides network-based drug repurposing results with network visualizations. AlzGPS will be a valuable resource to the AD research community. We will continue to add more types of omics data and update AlzGPS annually or when a large amount of new data is available. In summary, AlzGPS presents the first comprehensive *in silico* tool for human genome-informed precision medicine drug discovery for AD. From a translational perspective, if broadly applied, AlzGPS will offer a powerful tool for prioritizing biologically relevant targets and clinically relevant repurposed drug candidates for multi-omics-informed therapeutic discovery in AD and other neurodegenerative diseases.

## Declarations

### Ethics approval and consent to participate

Not applicable

### Consent for publication

Not applicable

### Availability of data and materials

All the data in AlzGPS can be freely accessed without registration requirement at https://alzgps.lerner.ccf.org.

## Authors’ contributions

F.C. conceived the study. Y.Z. constructed the database and developed the website. J.F., Y.Z., and Y.H.K. performed data gathering and processing. L.B., A.A.P., J.B.L., and J.C. discussed and interpreted all results. Y.Z. F.C., and J.C. wrote and all authors critically revised the manuscript and gave final approval.

## Acknowledgements

We thank the Lerner Research Institute Computing Services for hosting AlzGPS.

---

**Table S1.**
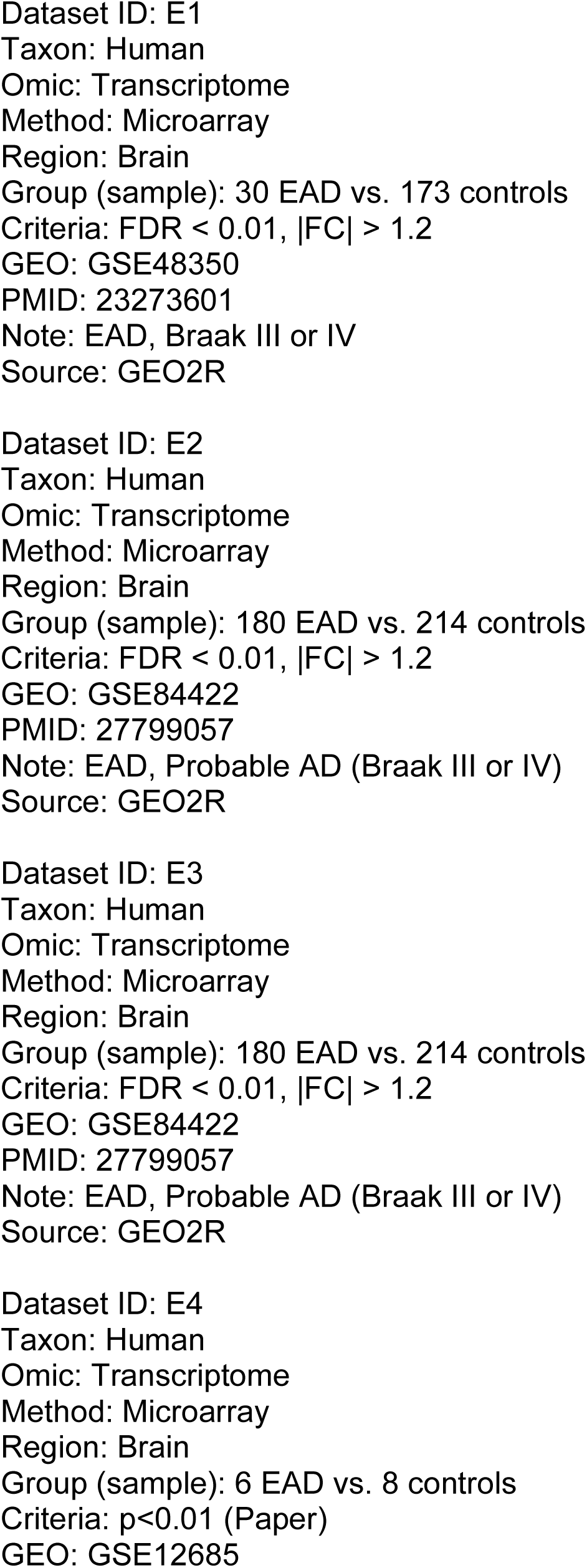
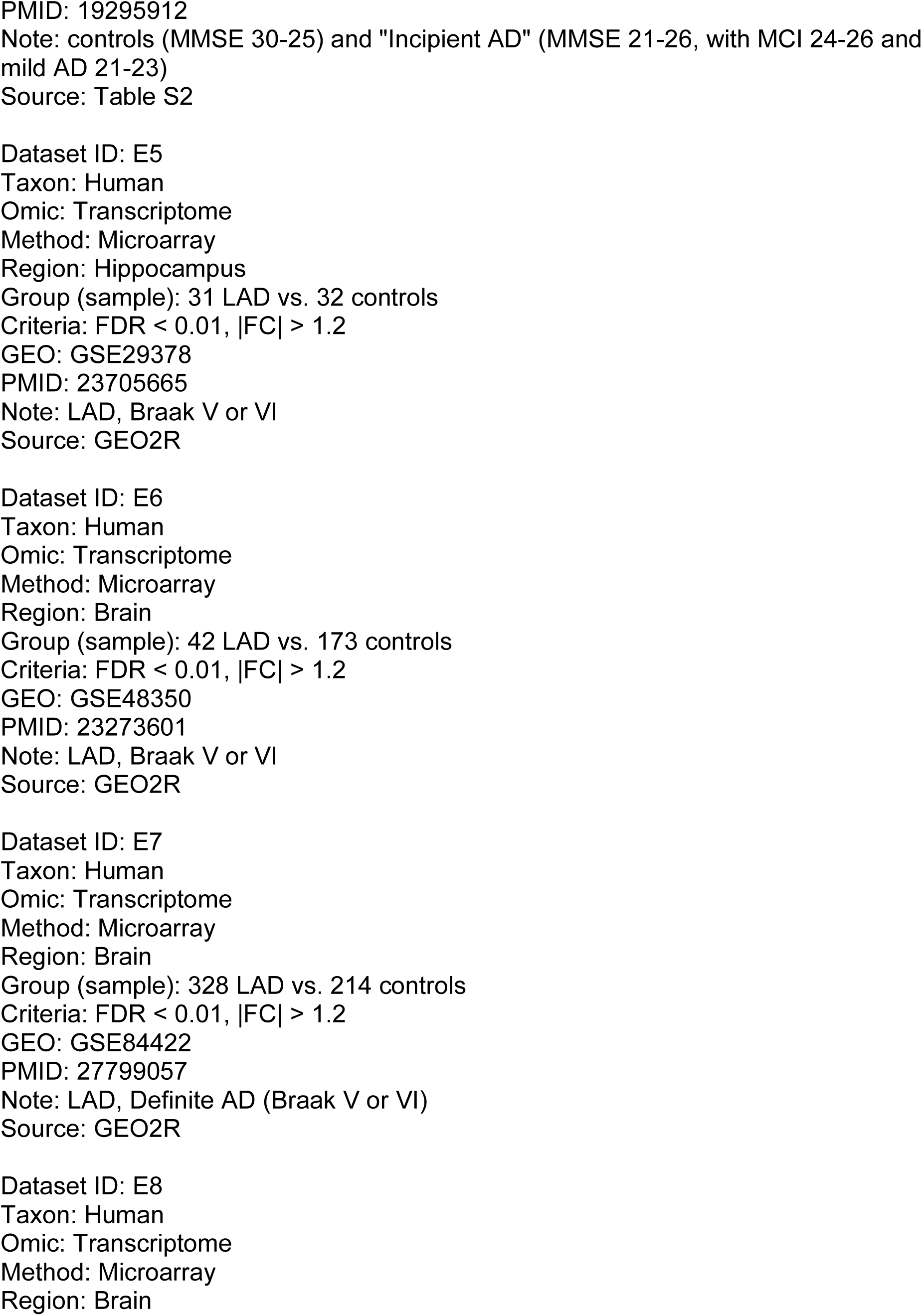
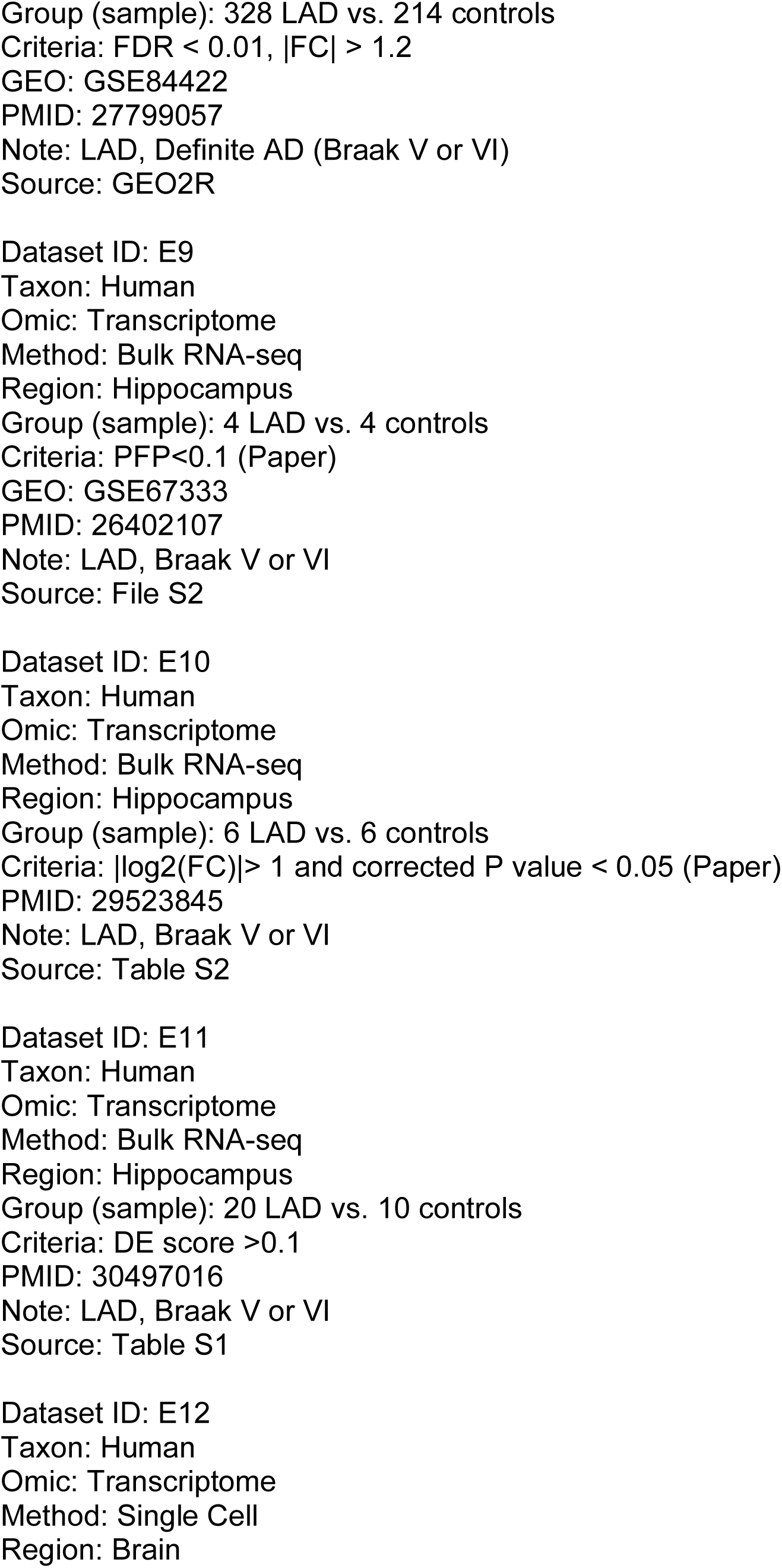
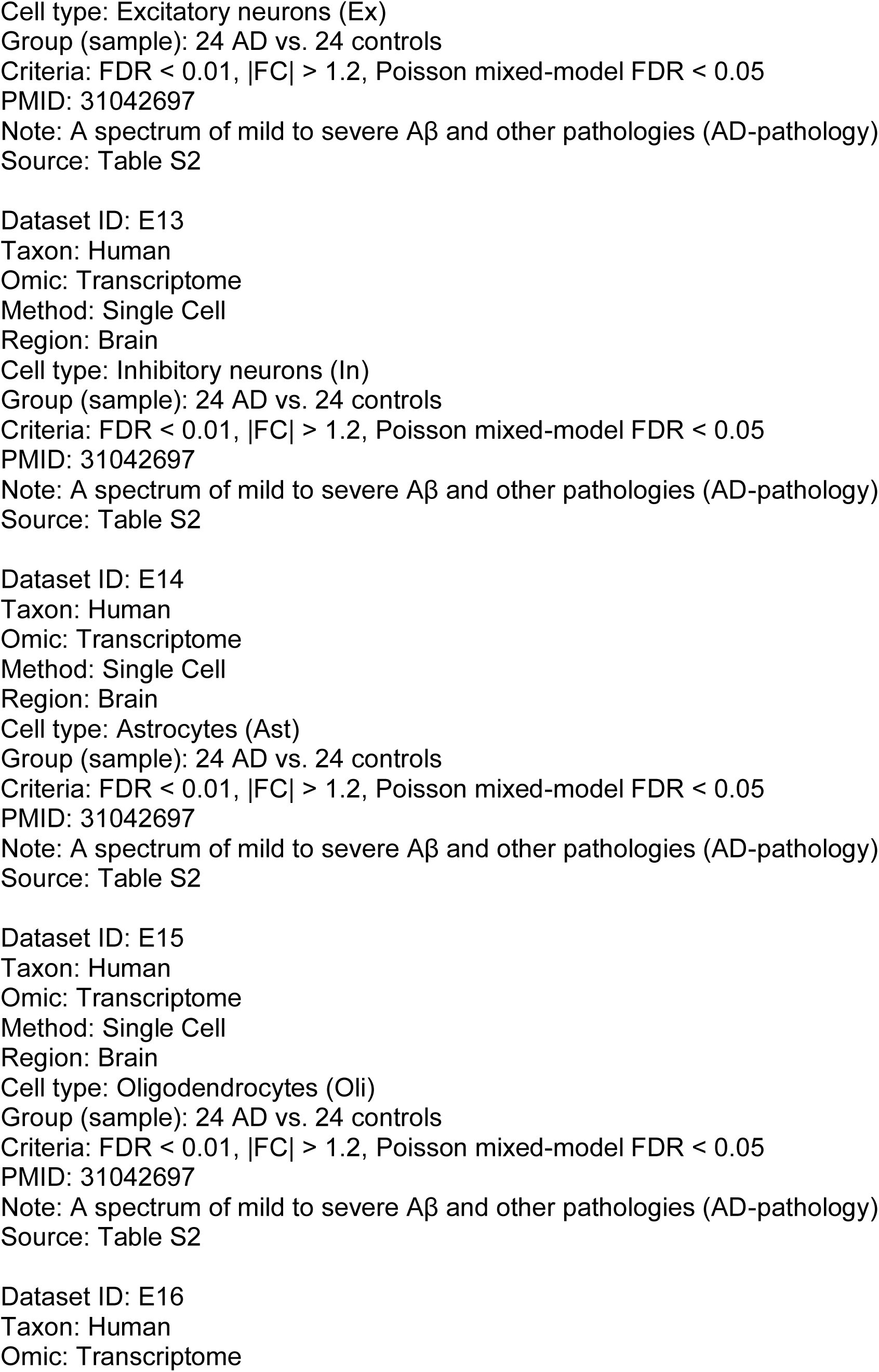
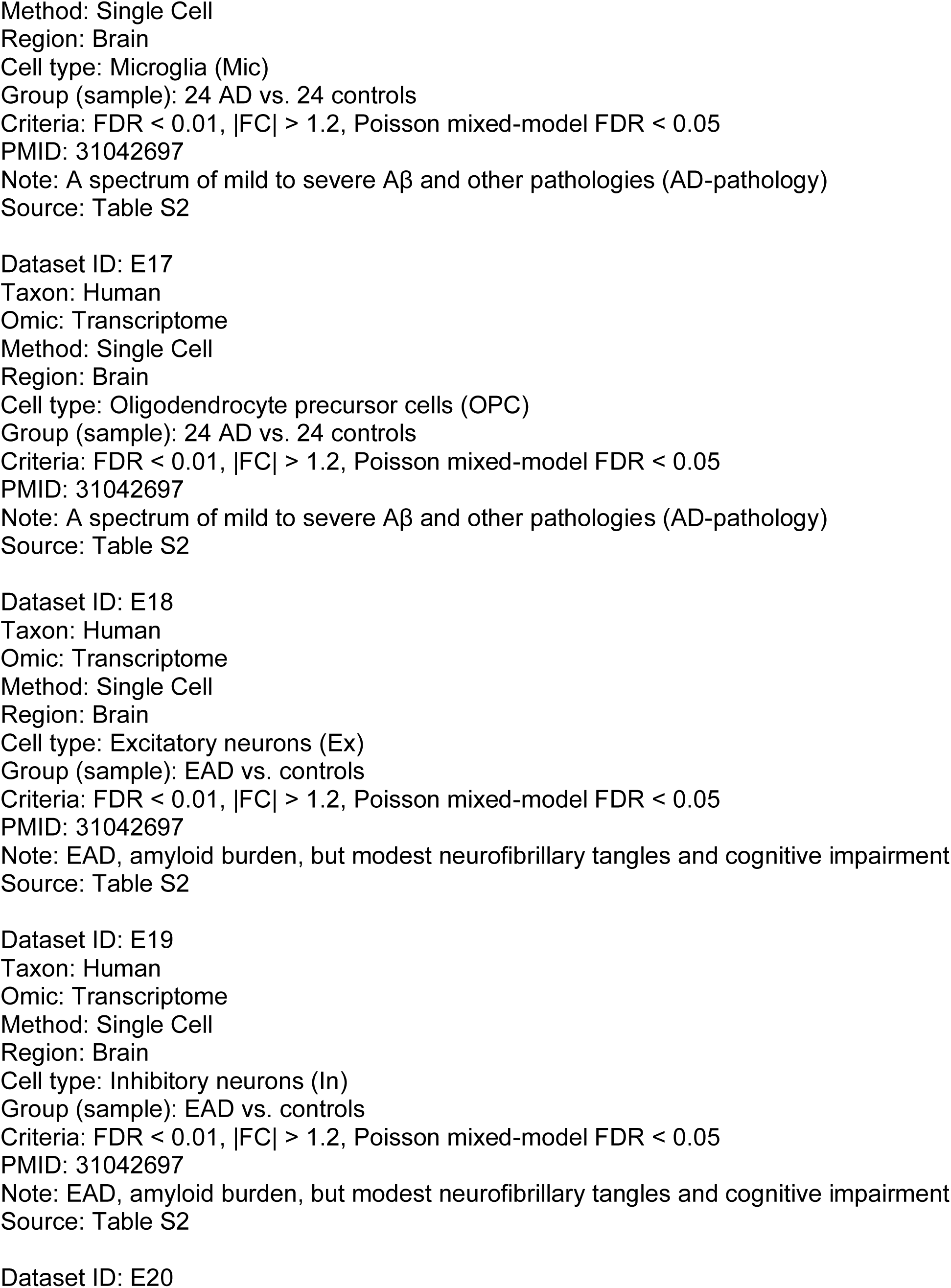
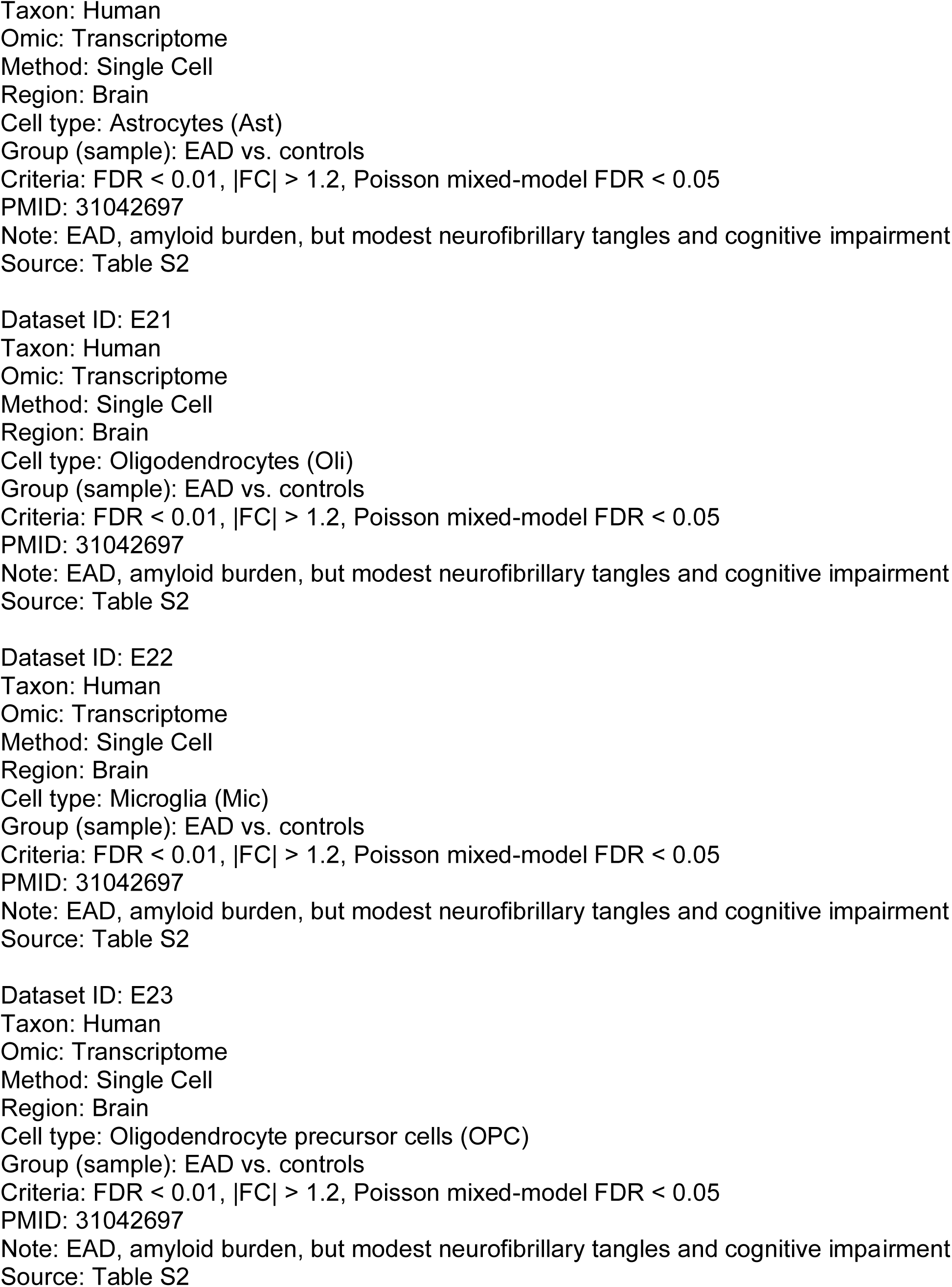
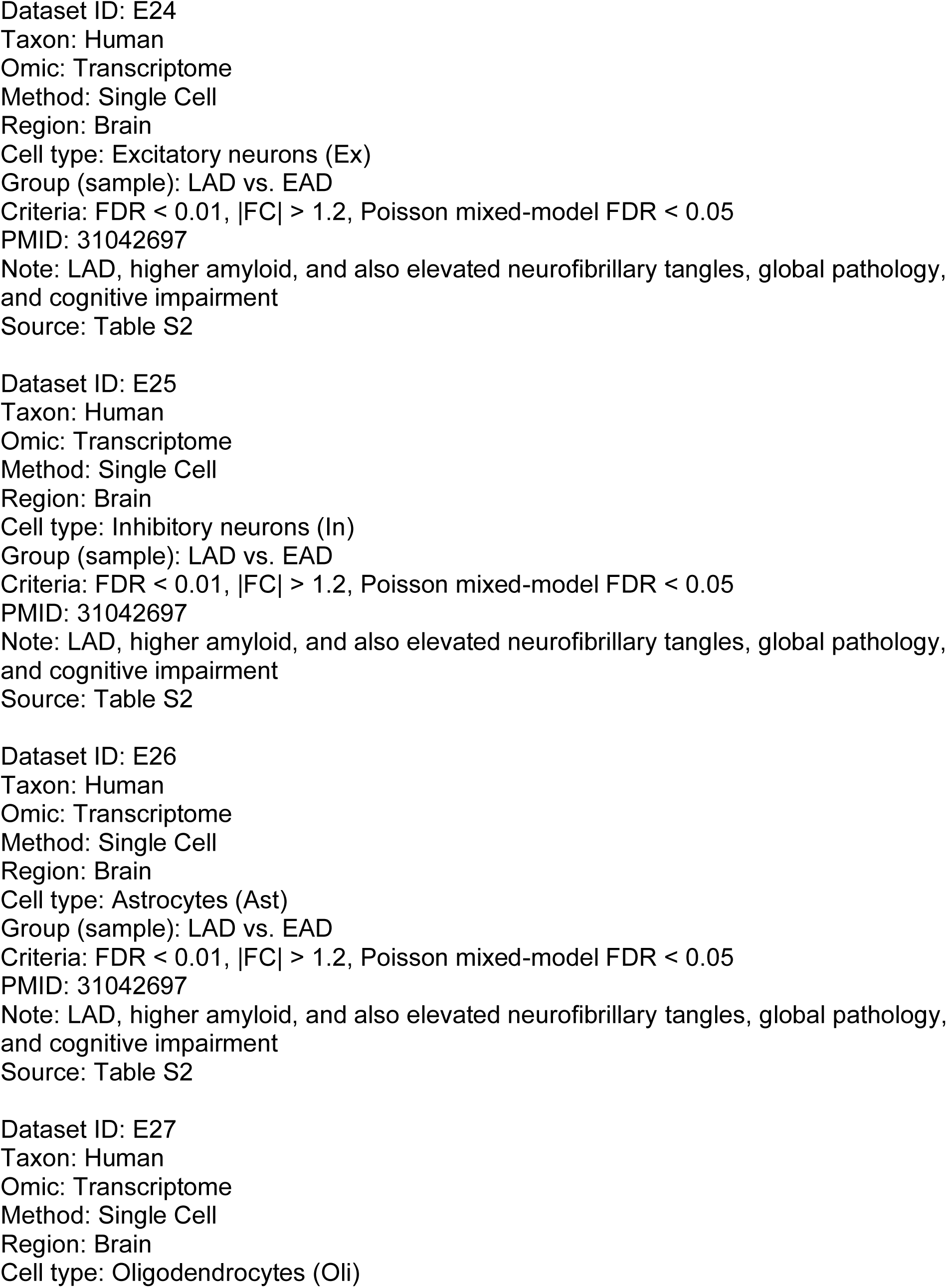
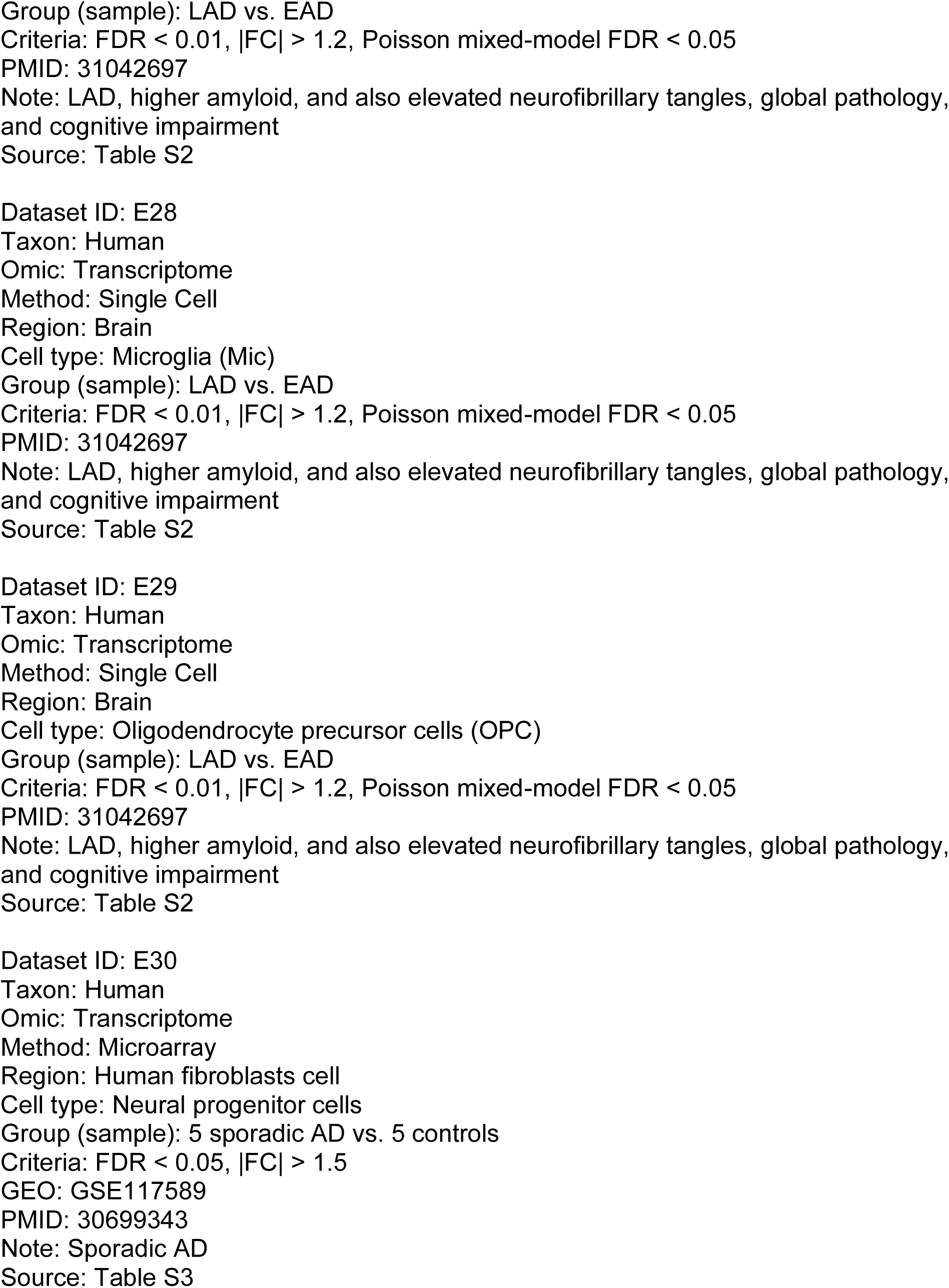
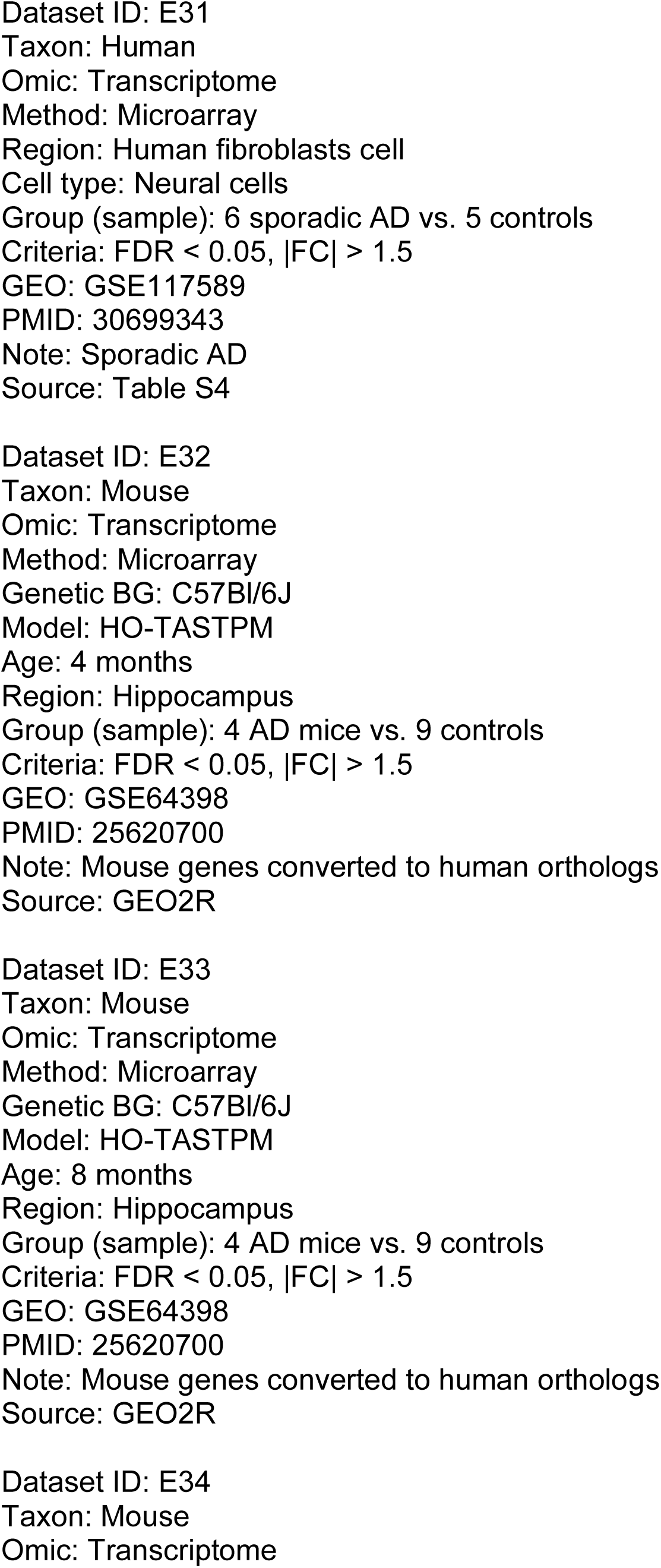
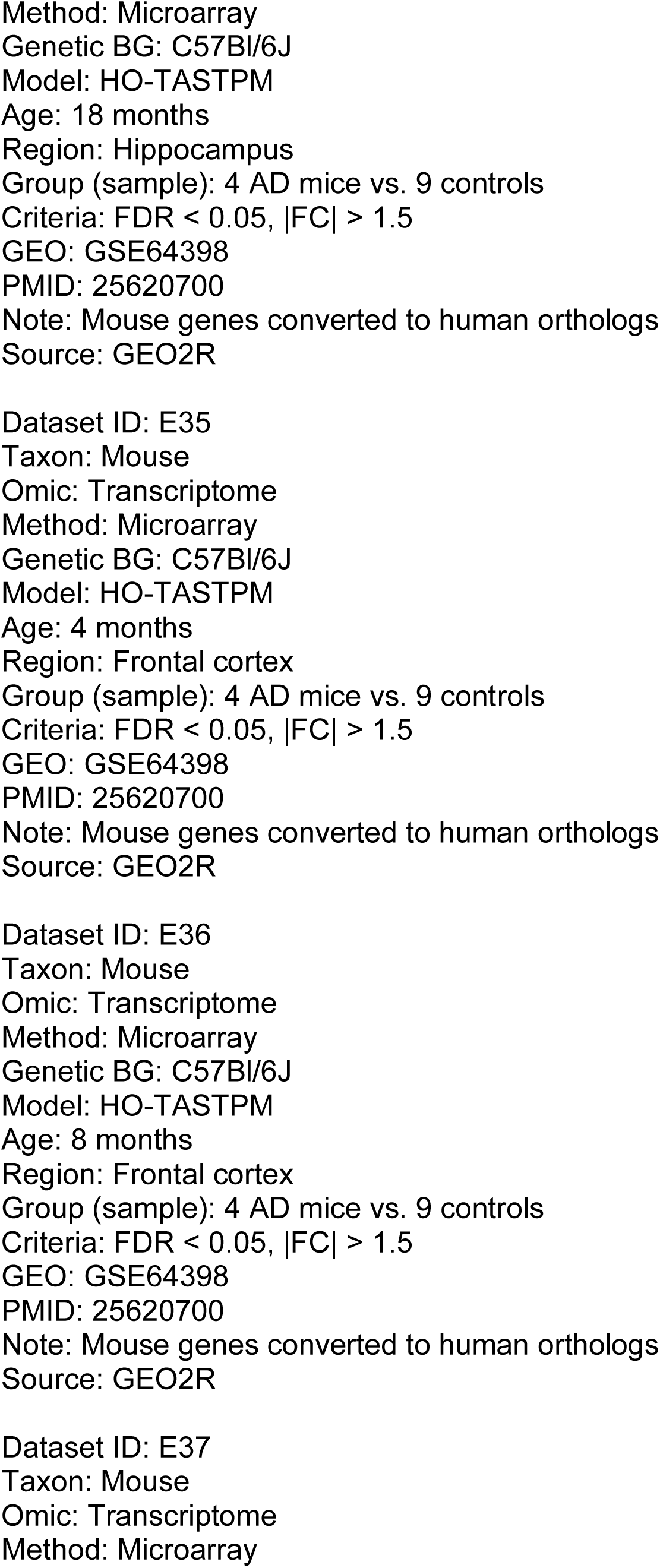
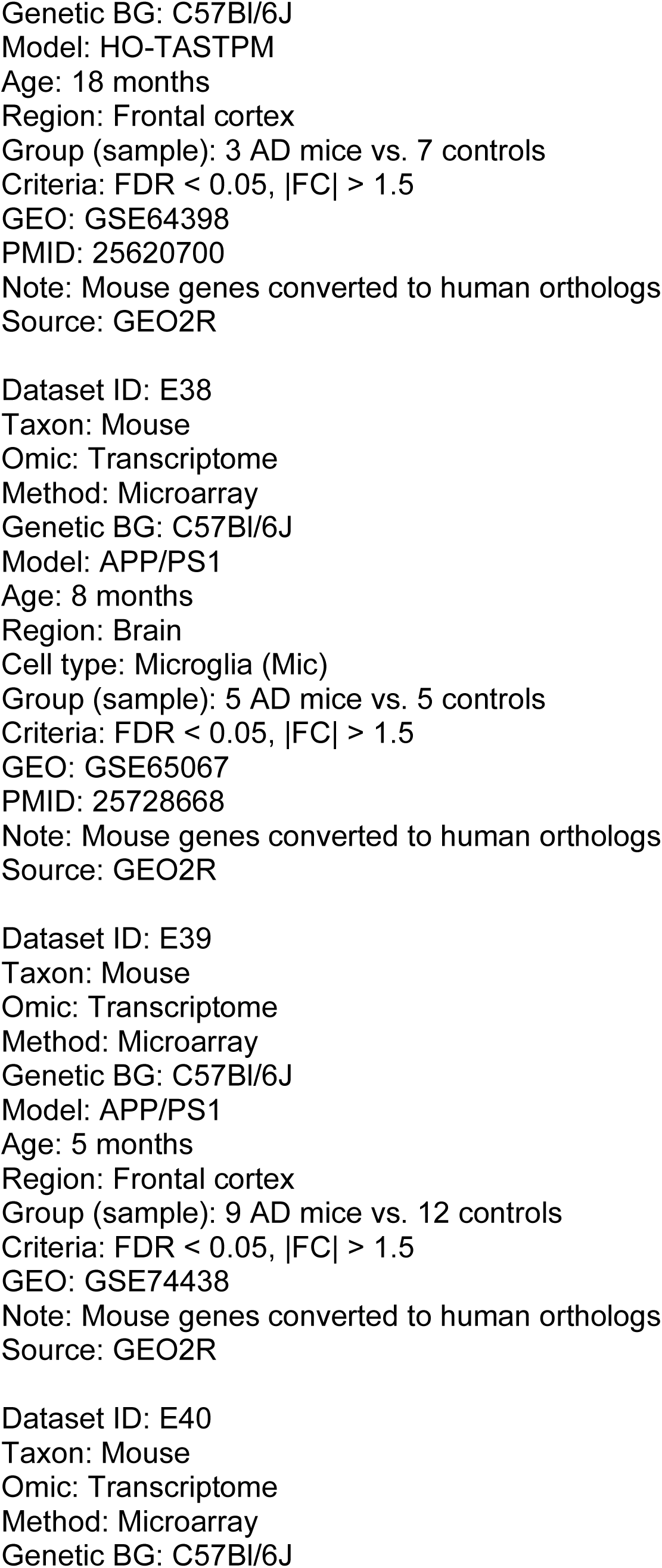
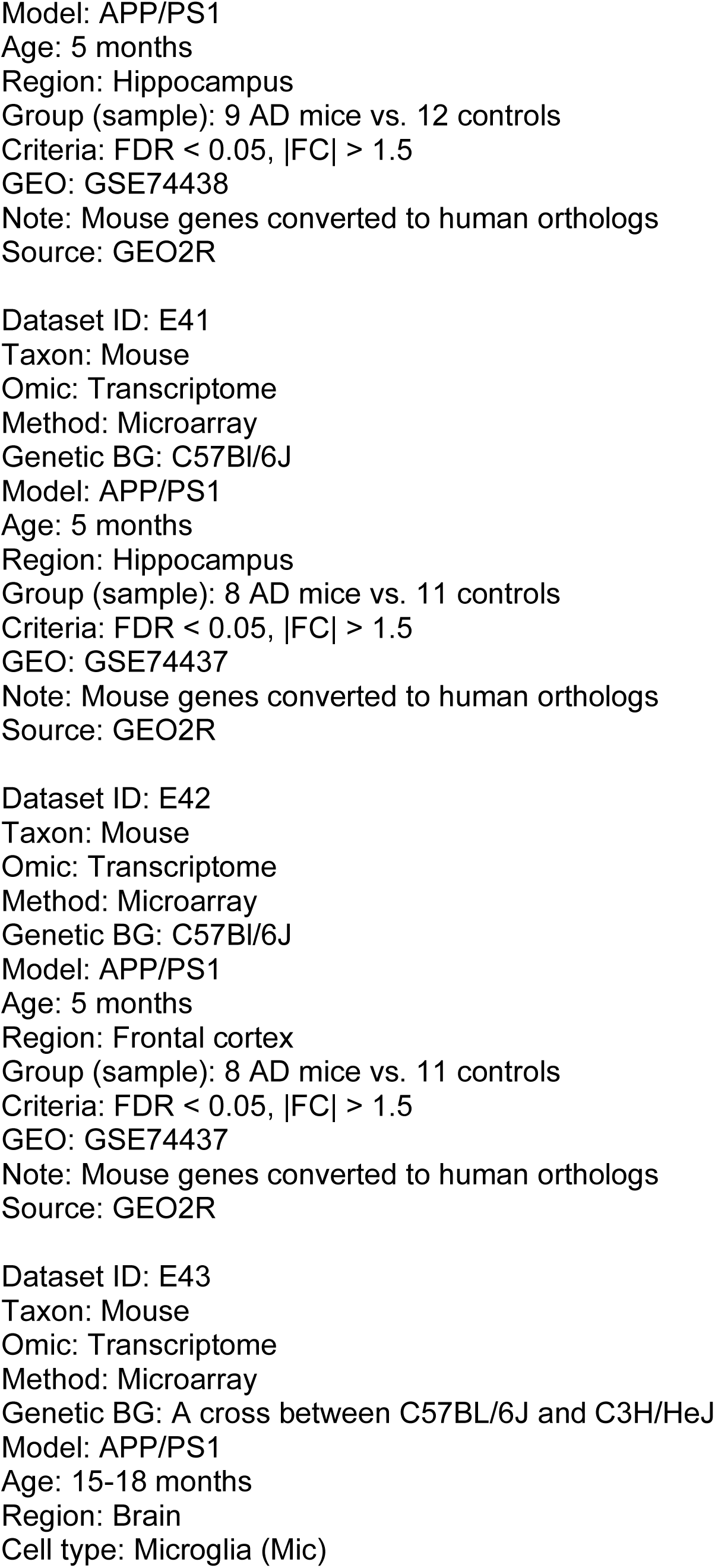
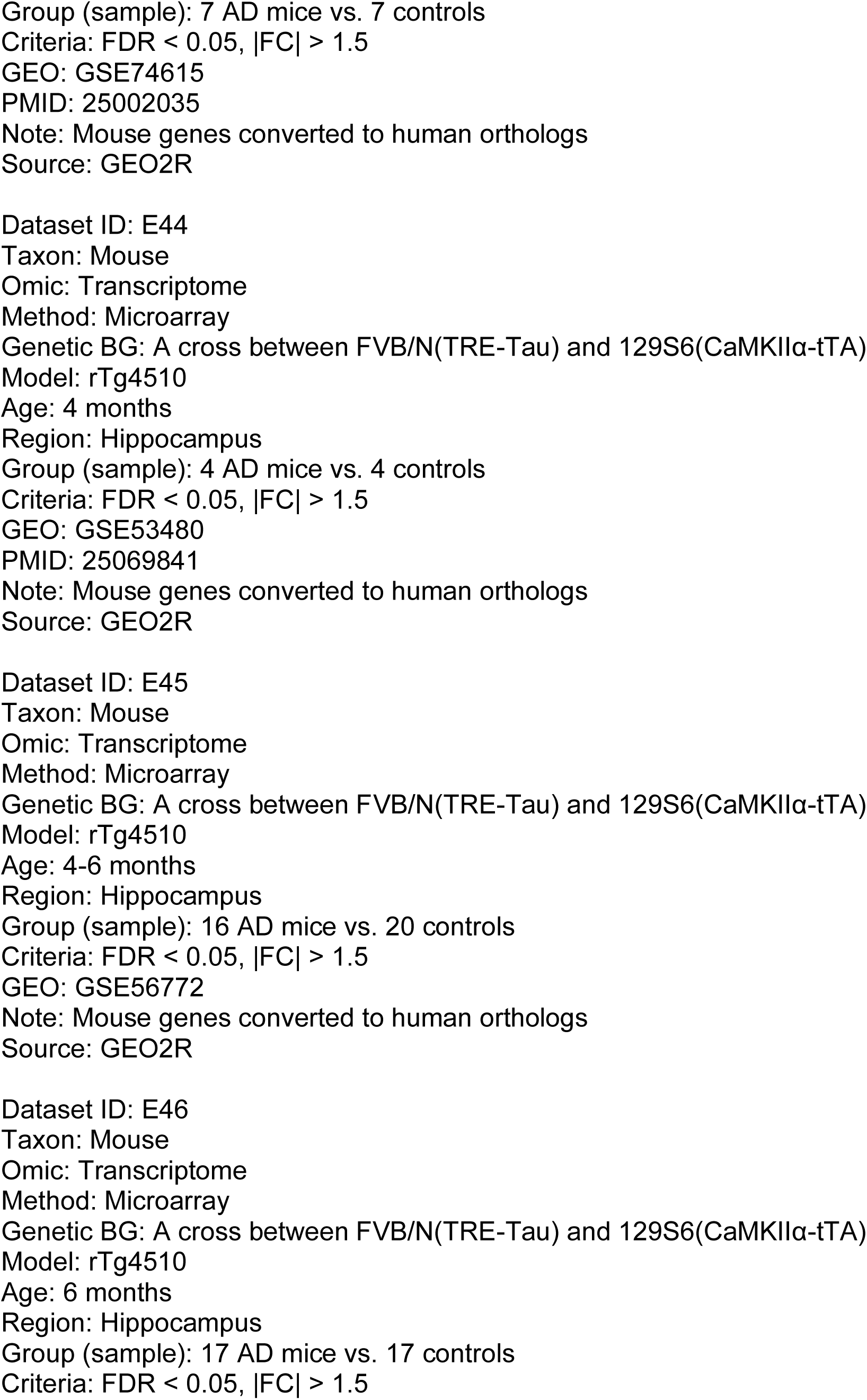
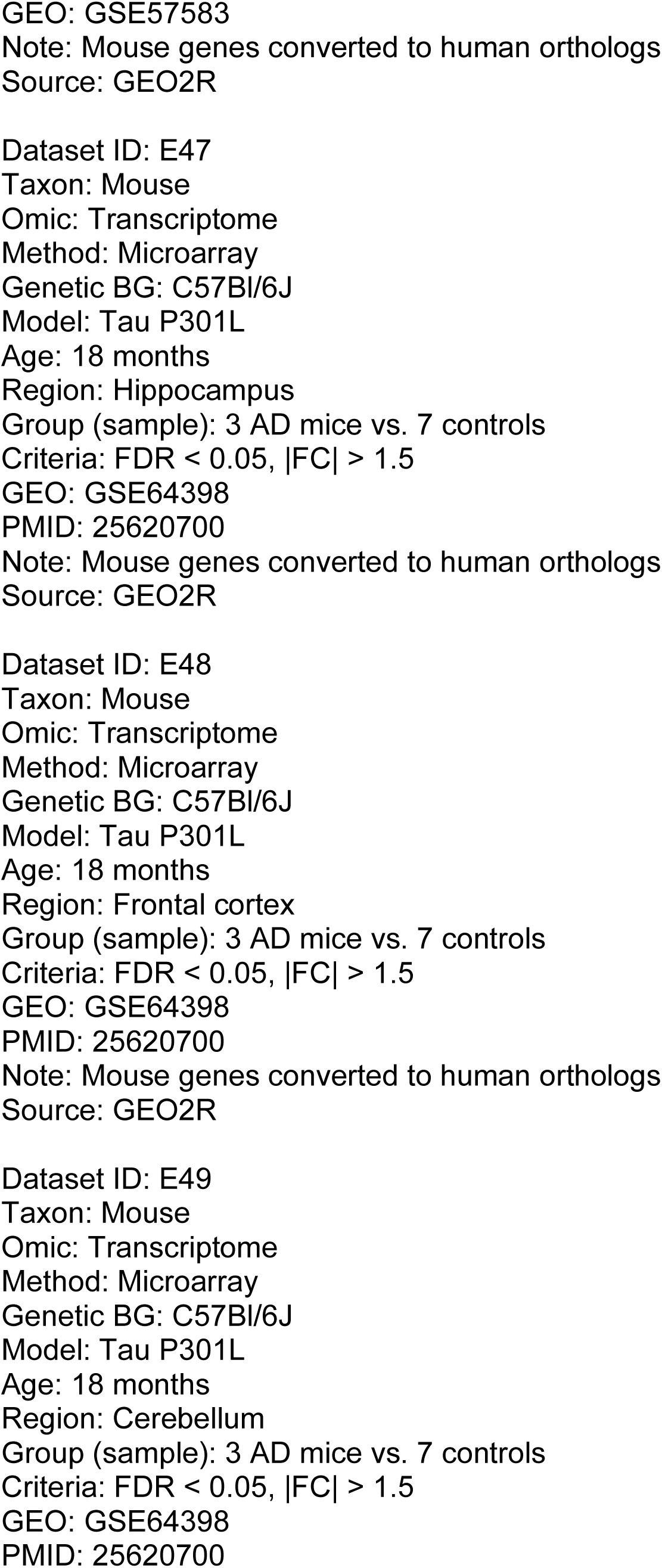
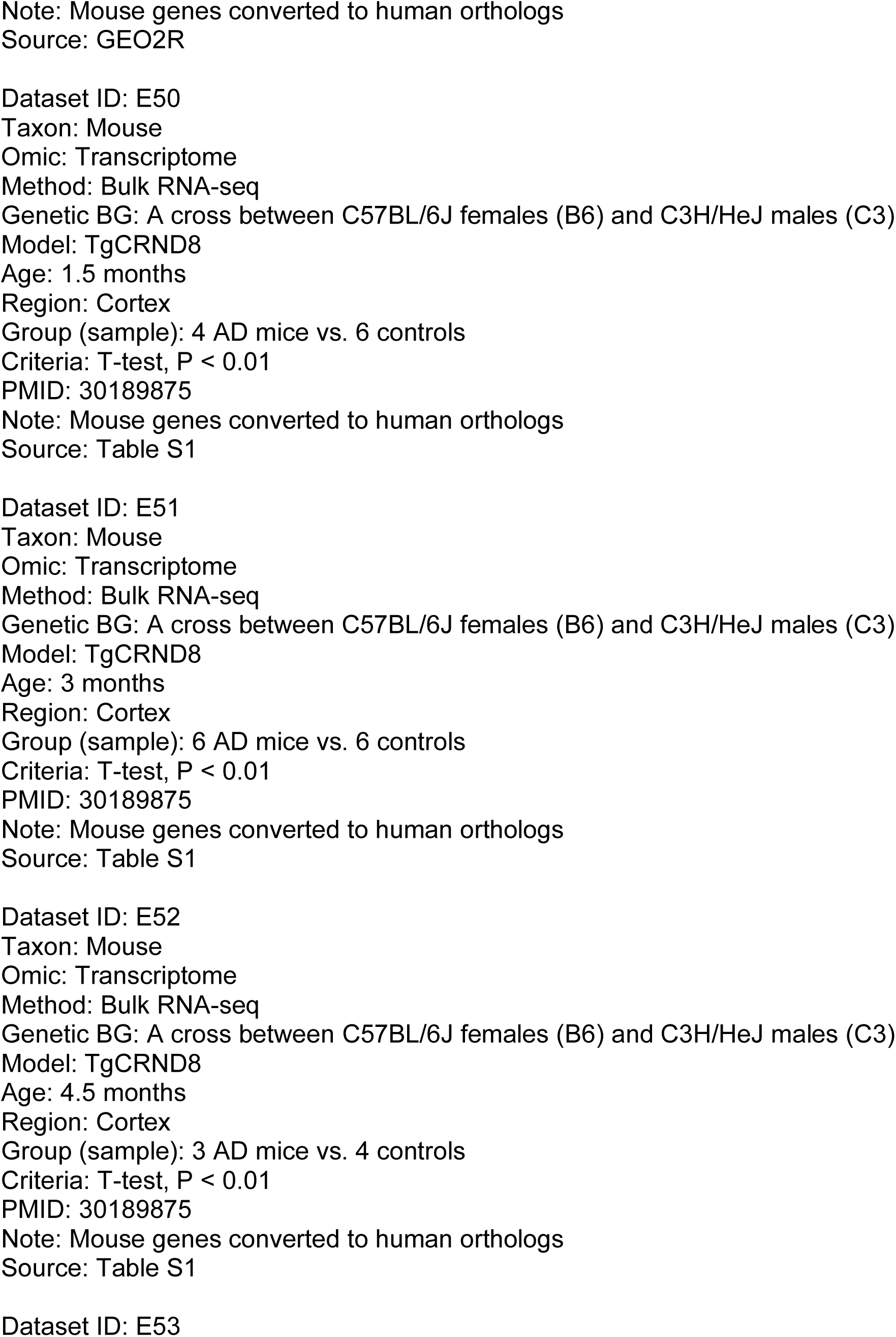
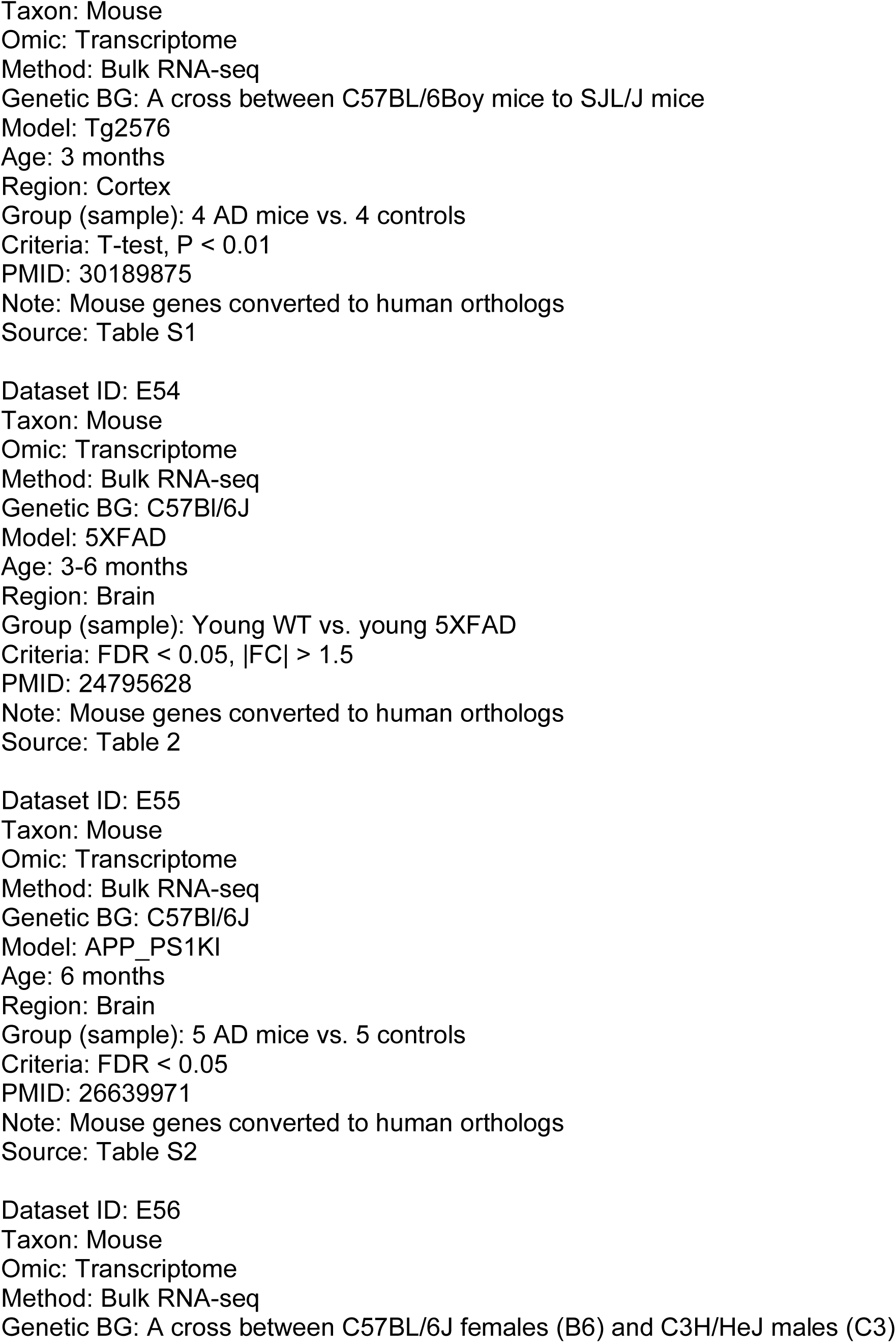
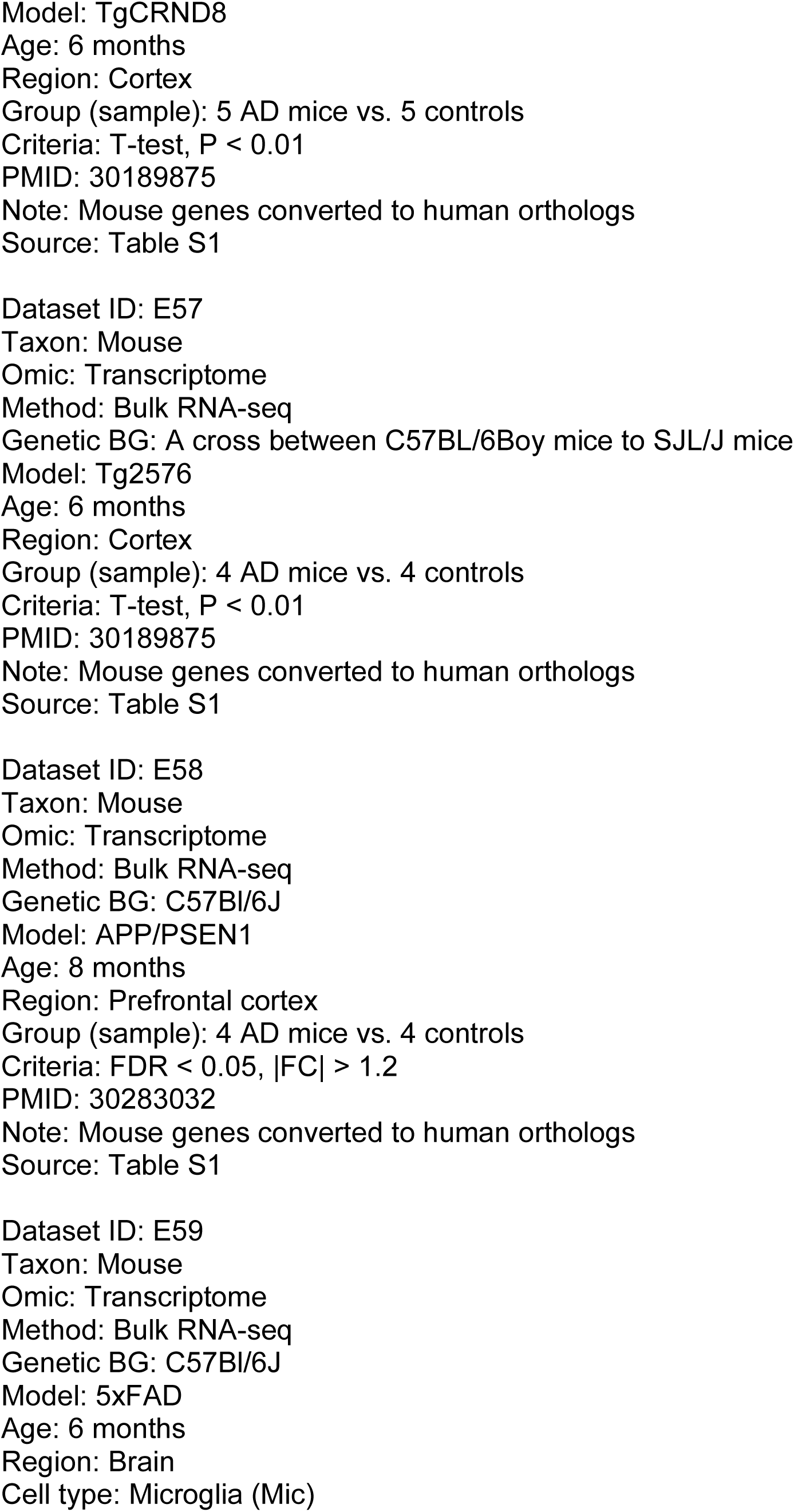
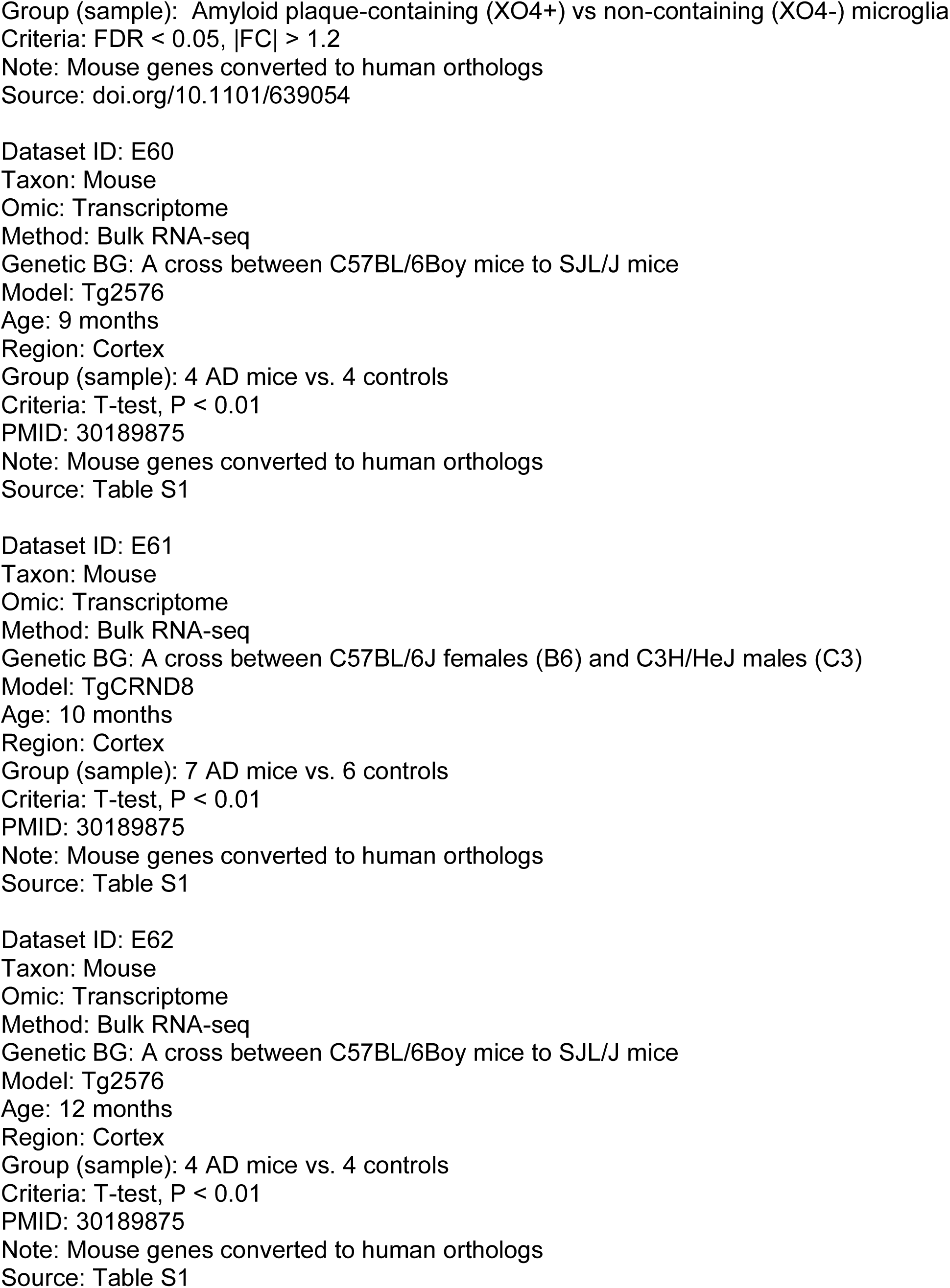
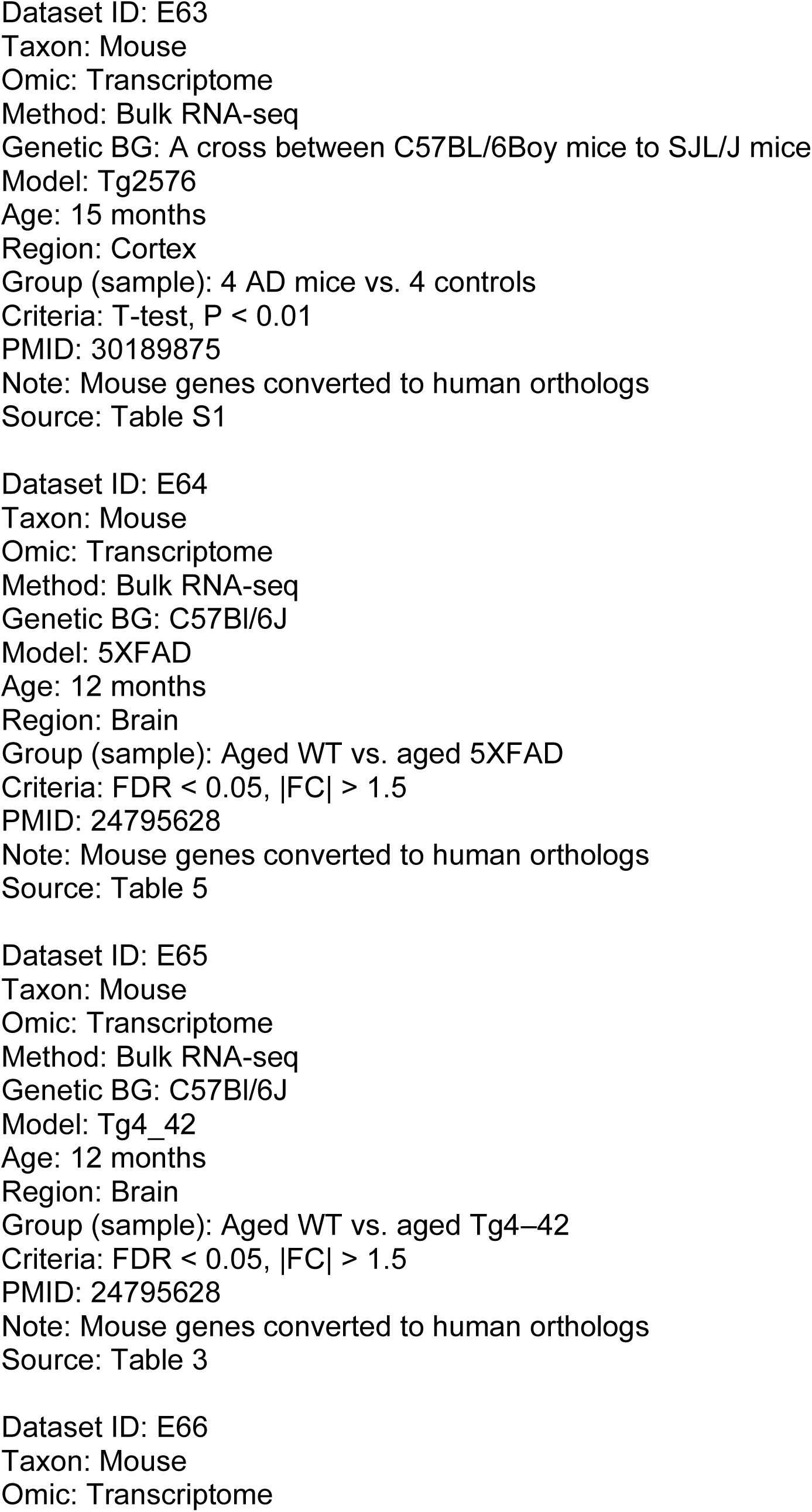
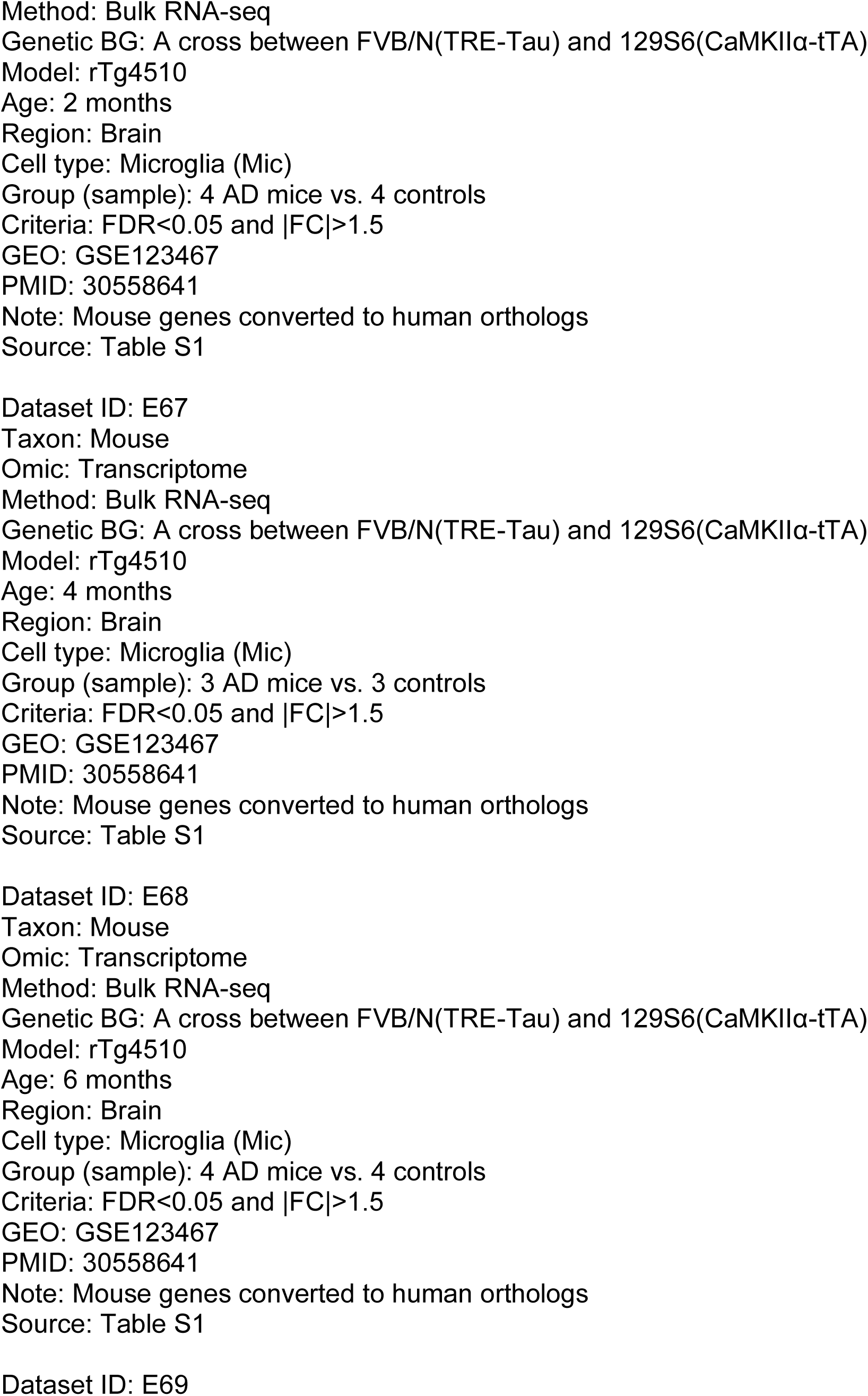
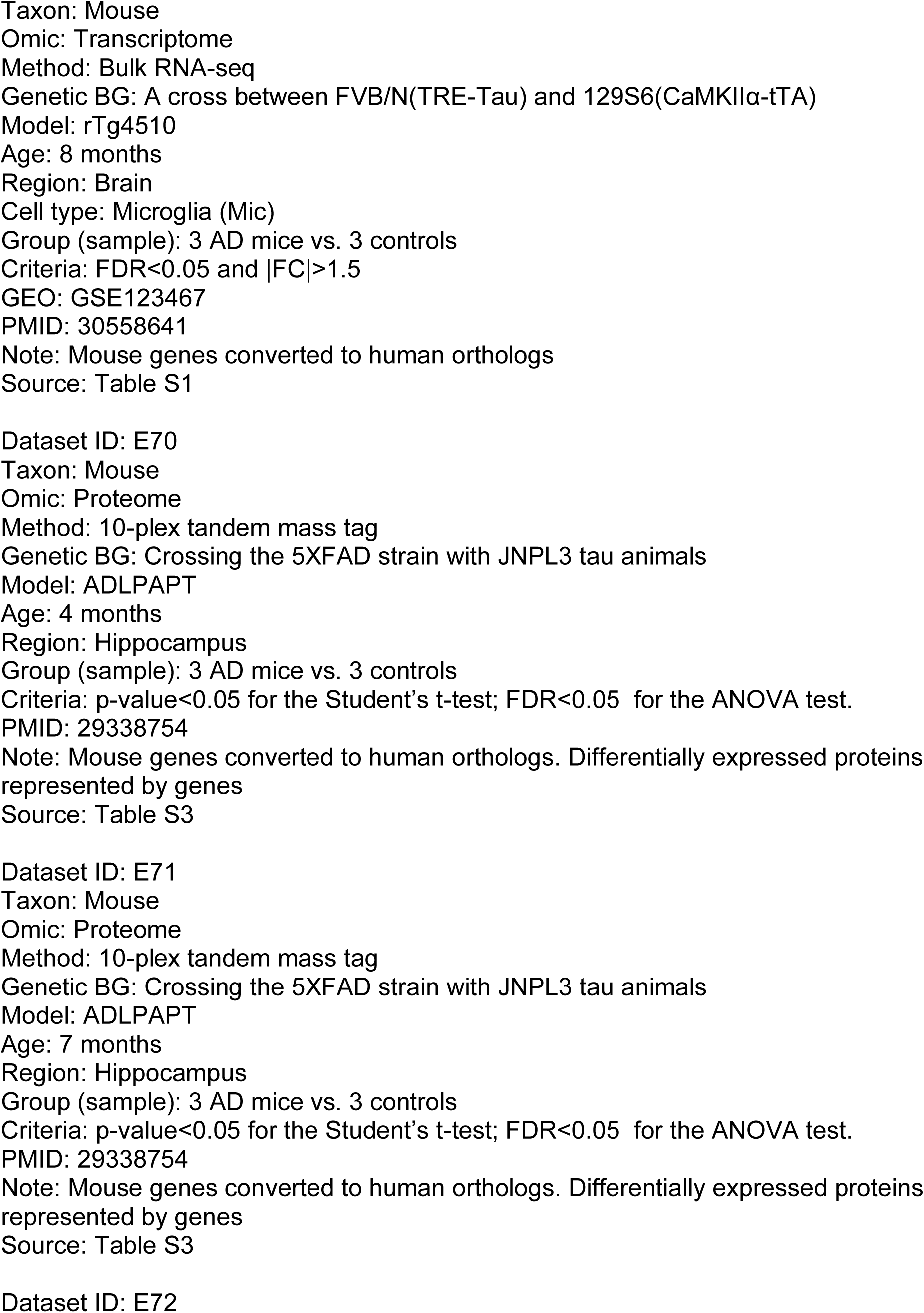
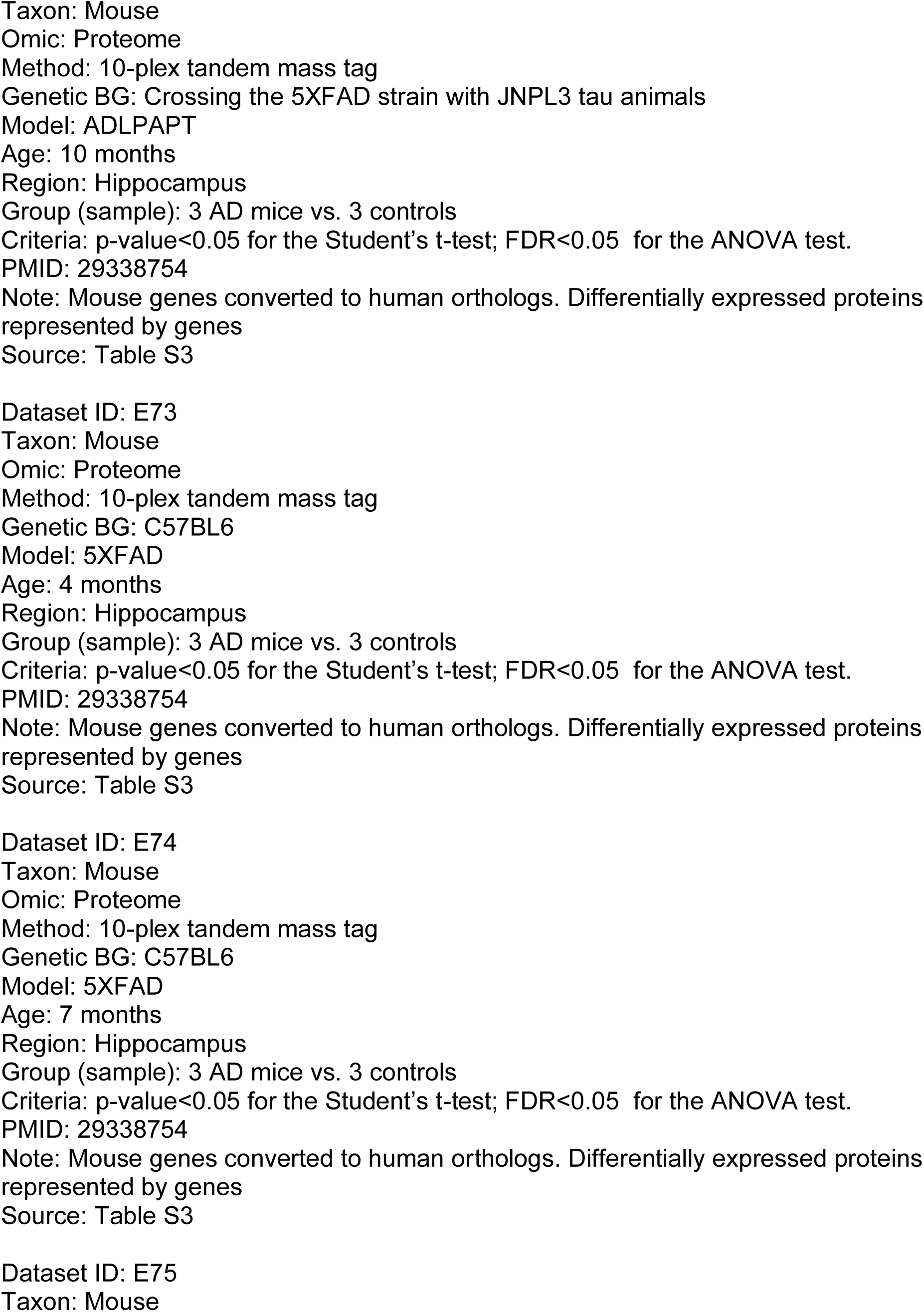
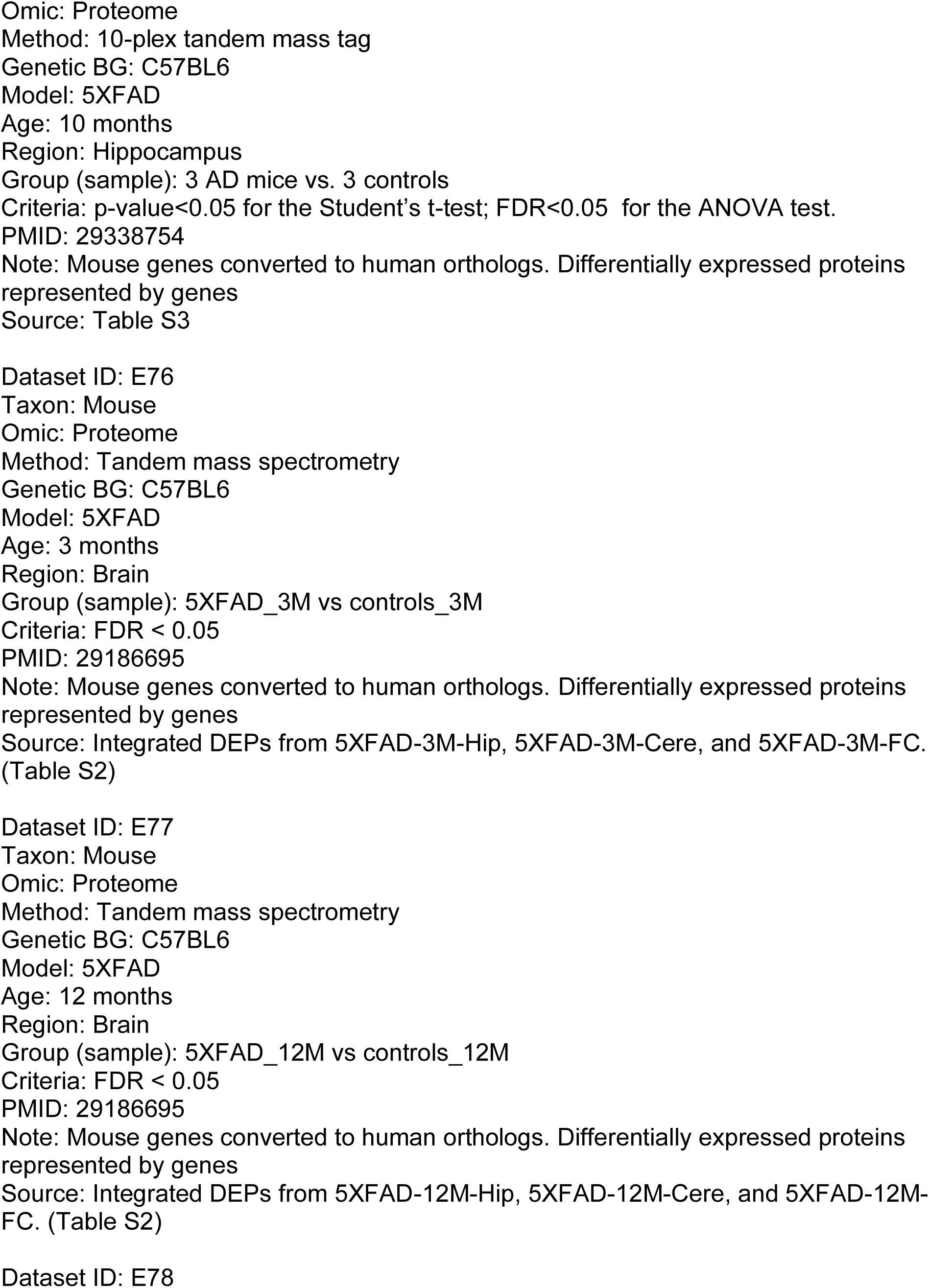
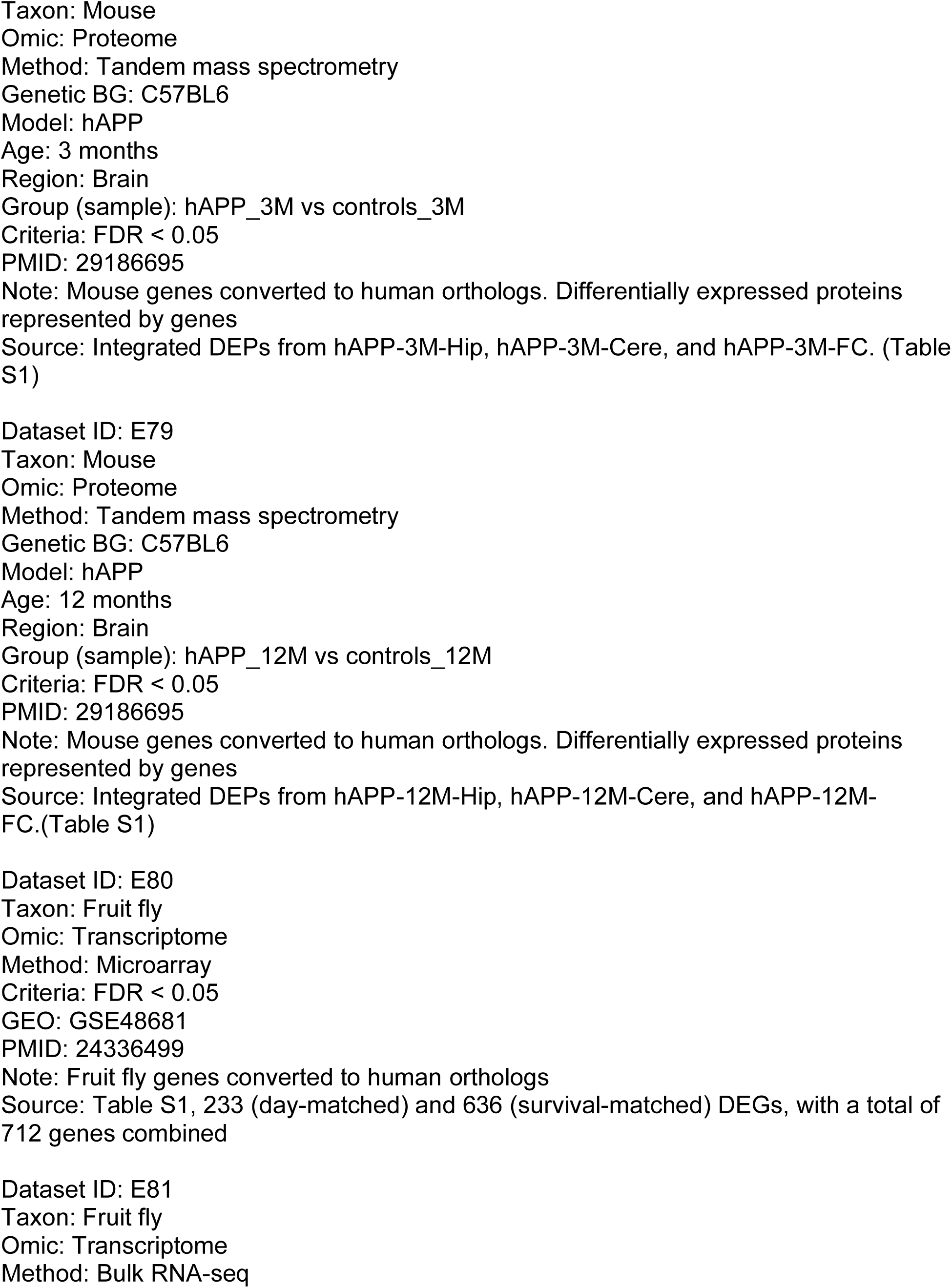
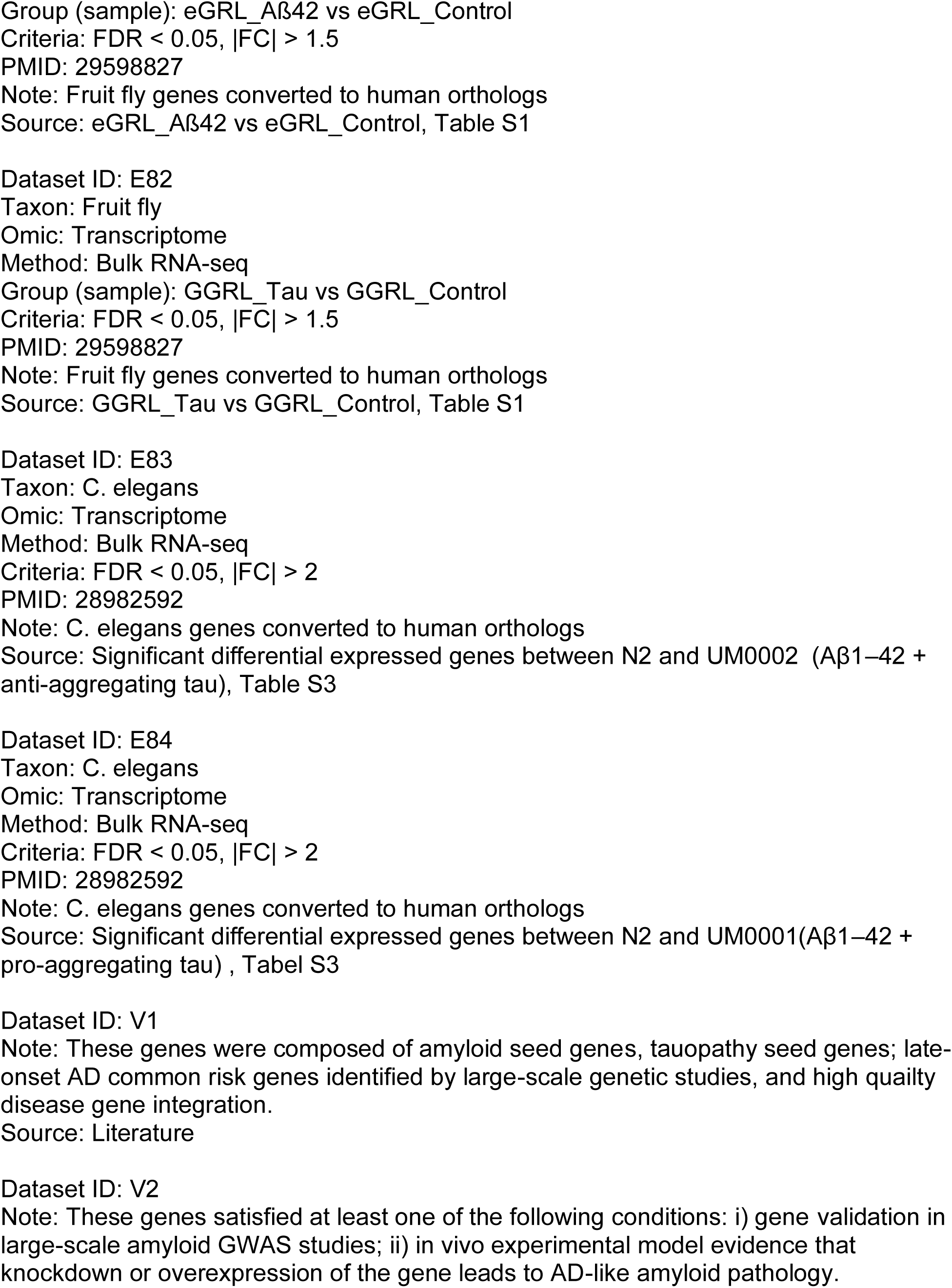
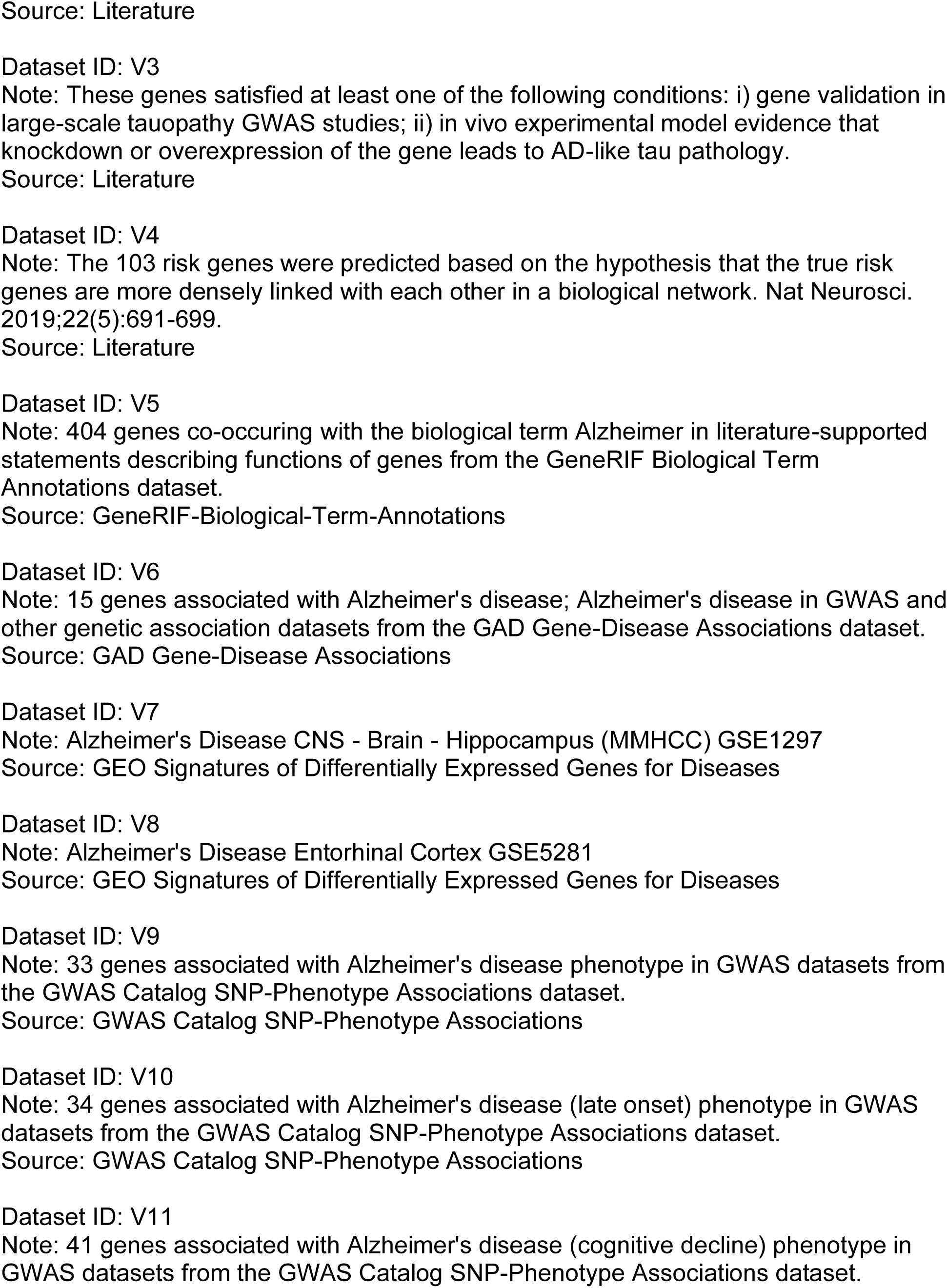
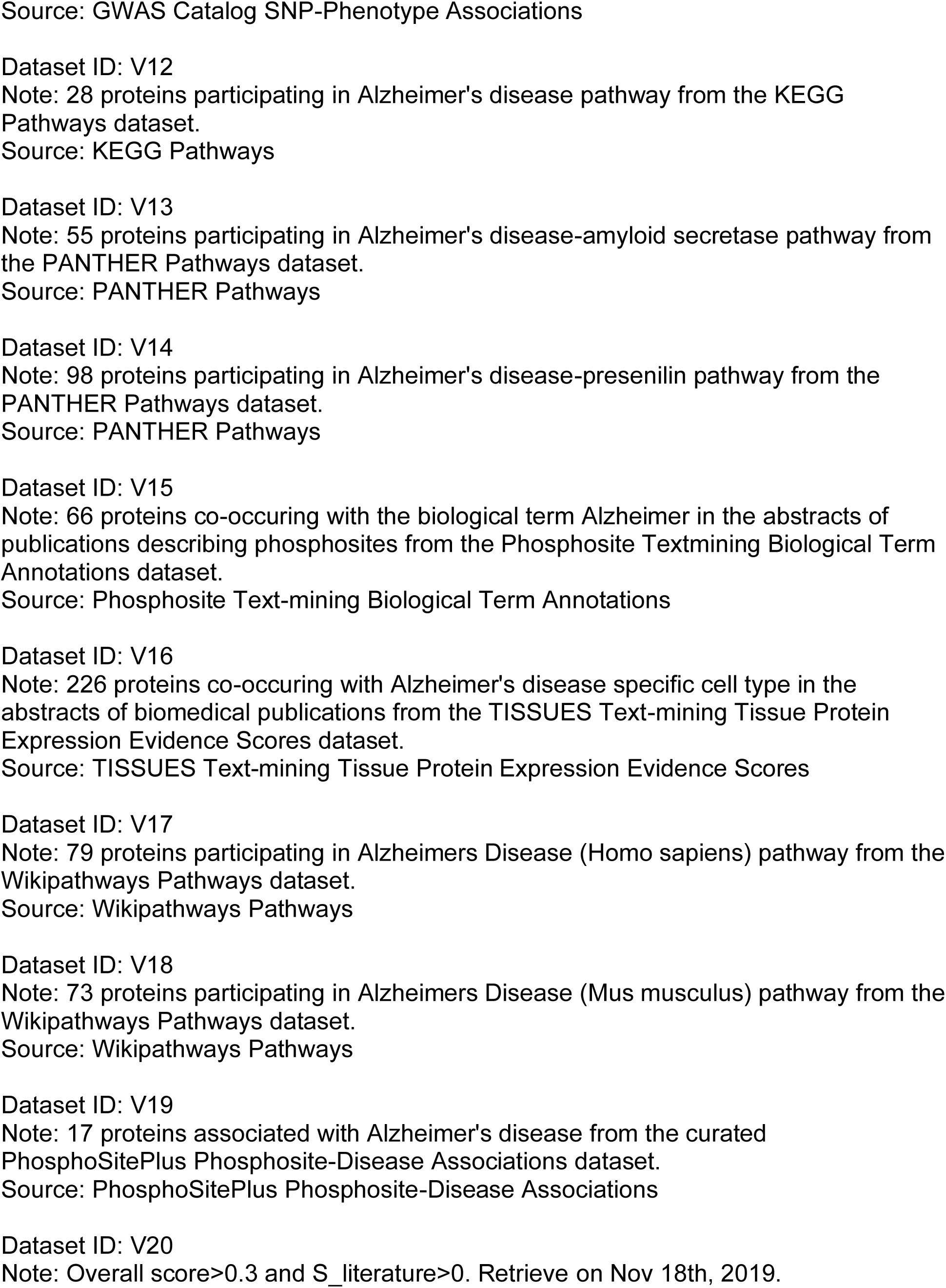
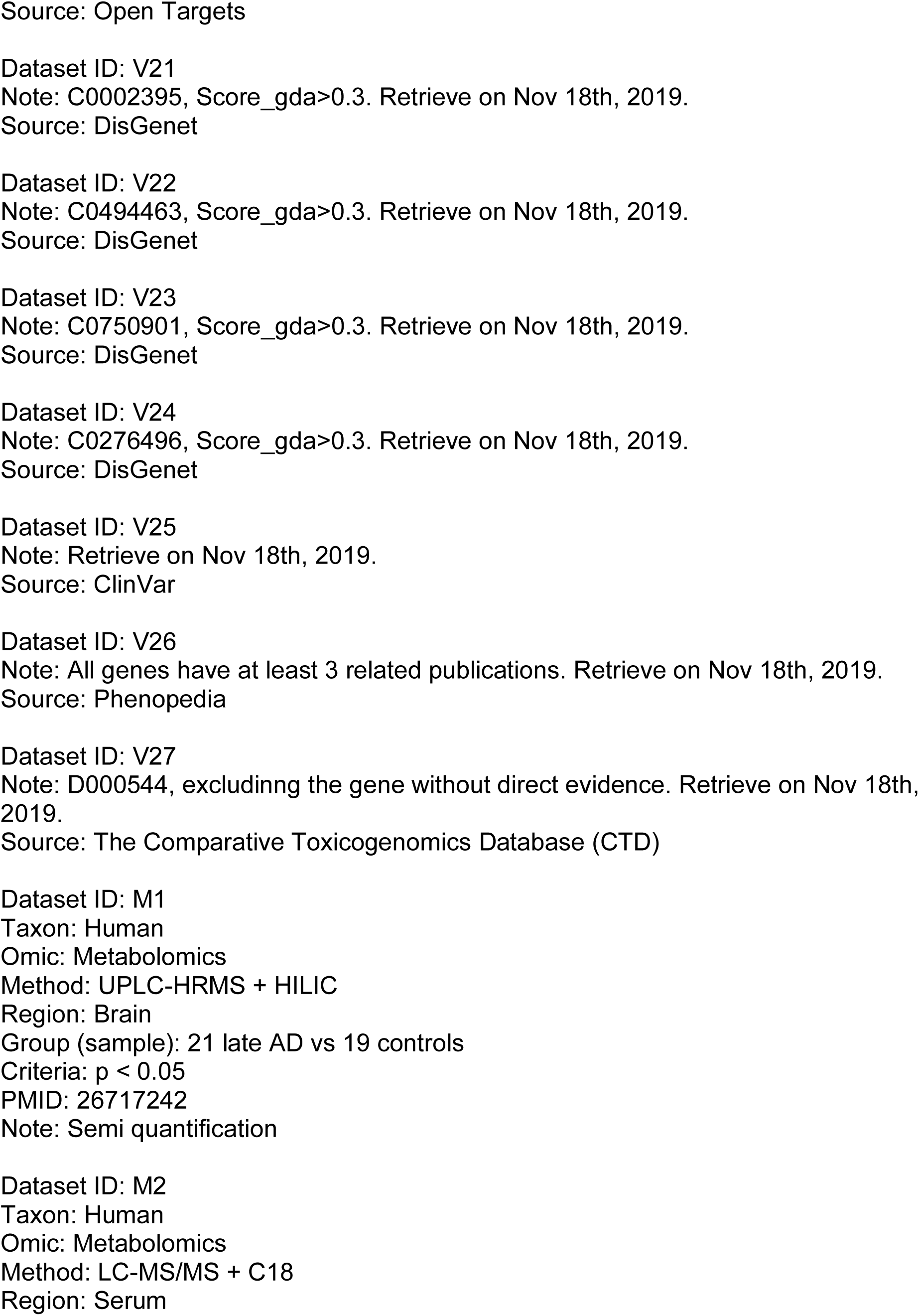
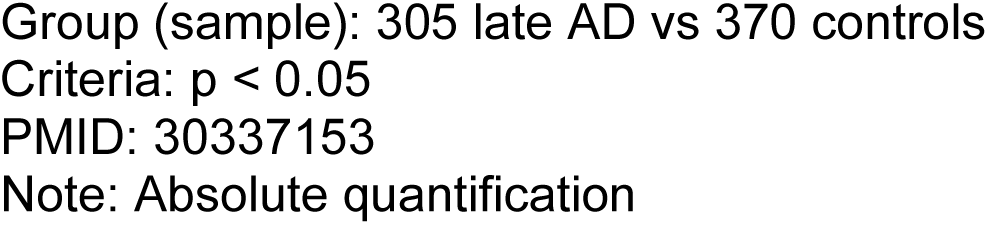
All data sets in AlzGPS.

